# Predicted and experimental protein-ligand coordinates with distance labels and evaluation splits

**DOI:** 10.64898/2026.09.06.749709

**Authors:** Thor Klamt

## Abstract

Structure prediction tools now provide protein–ligand geometry for pairs no laboratory experiment has resolved thus far. This data is typically provided without calibrated, practically usable per-system measures of whether a given complex is correct and how much to trust it. Agreement between independently constructed methods is an established signal for this type of problem. What has been missing is a way to derive what a given level of agreement is worth. We present PLI-Parallax, which deposits predicted geometry together with experimental data needed to calibrate it. Chai-1, two Boltz-2 configurations, and the docking engine smina were run over shared inputs across an experimentally resolved crystal tier of 19,350 complexes and a corpus tier of 31,746 cross-docked pairs without experimental ground truth on predicted receptors, yielding 307,314,646 residue-to-ligand-atom distance records, which the stored coordinates allow a consumer to recompute at a cutoff of their own choosing. Their mutual agreement is fitted against observed accuracy where experimental data permits it and carried to where it does not. This way 30,567 systems carry predicted label accuracy together with a split-conformal interval. On the crystal tier that interval covers observed accuracy at the stated rate. On the corpus tier it ranks systems by expected label quality, since both distributions differ. The deposit is accompanied by 646 evaluation configurations across seven data split families, and 631 of them report how far their training and test entities separate under a two-sample test. The 906 protein accessions were partitioned to control sequence leakage, so a protein-cold split here tests generalisation across sequence space. The two tiers support evaluation against experimental data and, where this is absent, supervision weighted by how far the configurations agree.

## Background and Summary

Binding outcomes are experimentally measured for far more protein–ligand pairs than for which experimentally resolved complex structures exist. Structure prediction closes that gap, and cofolding models and docking engines provide coordinates for pairs no laboratory experiment has yet resolved. Such tools commonly provide confidence scores alongside, and across several cofolding models those scores have been found not to separate correctly from incorrectly placed ligands [1]. Boltz-2 confidence values concentrate above 0.9 with essentially no low-confidence predictions, so a threshold on them separates little[2].

Several resources organise that geometry. PDBbind[3] pairs solved complexes with measured affinities. When it is reprocessed, a substantial share of the entries does not survive filters for covalent binding, rare elements, steric clashes and structures that cannot be repaired[4]. Others address what the record supports rather than what it contains, LP-PDBbind[5] and PLINDER[6] by controlling train–test similarity and MISATO[7] by adding molecular dynamics, so that the geometry becomes an ensemble rather than a single pose.

For pairs no crystal structure covers, resources now exist that provide generated coordinates from single teachers at scale with varying exclusion and selection strategies. SAIR[8] folds a million endpoint-labelled pairs with Boltz-1x[9], generating five samples per pair and selecting among them by agreement with an independent computational potency estimate rather than by the model’s own confidence. GatorAffinity-DB[10] folds several hundred thousand more with Boltz-1 itself, filtering on that model’s own confidence. The limits of that supervision are also documented. Runs N’ Poses[11] finds that cofolding methods largely reproduce poses they were trained on, most acutely for ligands seen in a single pocket. Predicted structures are nonetheless deposited as resources in their own right, at proteome scale[12] and for splice isoforms that no experimental structure covers[13]. For both SAIR and GatorAffinity-DB the criterion for keeping a prediction comes from within the system that produces it.

Each element of the design that follows has a precedent, and Table 1 sets out where this deposit sits among them. Scale is not where it differs, since SAIR[8] and GatorAffinity-DB[10] carry roughly twenty and nine times its 51,096 systems. Runs N’ Poses[11] runs several cofolders over shared inputs and publishes the coordinates with training-similarity metadata, which is the nearest precedent for the construction used here. SPECTRA[14] generates splits across a spectrum of cross-split overlap and reads a model’s performance along it, which requires training a model and is not attempted here. CalPro[15] and reliability-aware multi-engine fusion[16] both estimate reliability from a model rather than depositing it as a field.

**Table 1.** Scope of this deposit against related protein–ligand resources. The axes are those on which the resources differ. Each count is the one its own release reports. Geometry names what produced the deposited coordinates. Anchor records whether the artifact holds predicted protein–ligand complexes together with measured complexes against which those predictions can be scored. PDBbind and MISATO hold no predicted complexes, MISATO’s molecular dynamics being simulated conformations of measured ones. PLINDER links AlphaFold receptor models to its holo systems but does not deposit predicted complexes. CrossDocked2020 holds cognate poses alongside its non-cognate ones. Calibrated records whether a per-system estimate of label quality ships with the data. The estimate must have been fitted against observed accuracy on measured complexes. Several of these resources ship a model’s own confidence score or a physical-validity flag, neither of which has been fitted in that way.

| Resource | Scale | Geometry | Anchor | Calibrated | Splits |
| --- | --- | --- | --- | --- | --- |
| PDBbind[3] | 19,443 (v2020) | experiment | no | no | none |
| MISATO[7] | 16,972, PDB-bind entries | experiment, MD | no | no | none |
| PLINDER[6] | 449,383 | experiment | no | no | leakage-controlled |
| CrossDocked2020[17] | 22.5M poses | docking | partial | no | clustered CV |
| SAIR[8] | 1,048,857 | cofolding | no | no | none |
| GatorAffinity-DB[10] | 456,526 | cofolding | no | no | none |
| Runs N’ Poses[11] | 2,600 | cofolding | yes | no | post-cutoff, similarity-graded |
| PLI-Parallax | 51,096 | experiment, cofolding, docking | yes | yes | 646 across seven families |

Agreement between methods of different construction is an older signal than any of those, and for classical docking it was a strong one. Requiring two programs to agree raised pose prediction from 55, 58 and 64 % for single programs to 82 % or above[18]. What has been missing is a way to say what a given level of agreement is worth, which requires cases where the answer is known. This deposit computes that signal over cofolding models and a docking engine, calibrates it against experimental structures where those exist, and releases the result as a per-system field.

PLI-Parallax is a resource of predicted and experimental protein–ligand coordinates with distance labels derived from them. Each system stores three-dimensional coordinates for receptor and ligand from four teacher configurations run over shared inputs, so that ligand-to-residue, residue-to-residue and ligand-to-ligand distance maps could be derived by the consumer rather than fixed by the deposit. They are neither repeated sampling of one method nor four independent ones. Chai-1[19] and Boltz-2[20] both follow the AlphaFold3[21] architecture and draw their training data from the same structural archive, with Boltz-2 contributing two configurations, one single-sequence and one alignment-conditioned. The physics-based docking engine smina[22] follows neither, optimising a scoring function rather than learning from structures. Three of the four configurations thus come from one structure prediction lineage.

The corpus tier holds 31,746 cross-docked pairs modelled on AlphaFold[23, 24] receptors, where no experimental complex exists for the pair, while the crystal tier holds 19,350 experimental complexes selected from BioLiP2[25] and the Protein Data Bank[26], against which the predicted coordinates can be measured. Across both the deposit carries 307,314,646 residue-to-ligand-atom distance records over 51,096 systems. Counts vary by denominator throughout, falling as the requirement defining each denominator tightens. Systems covered by all three of Chai-1, Boltz-2 single-sequence and the docking arm carry a predicted label accuracy with a split-conformal interval, released as a field for 30,567 of them. 646 train, validation and test configurations are provided across seven families, five of which span both tiers. Each configuration is redundancy-reduced at a stated maximum rather than leakage-free.

The deposit trades volume for a tier where the answer is known. Its coordinates and the distance labels derived from them are the substrate, and the reliability field and the split family are built on top of them. The two tiers are used differently. On the crystal tier the predictions can be scored against experiment, so the resource records what each configuration produces under throughput-oriented settings. On the corpus tier no such scoring is possible, and the labels are geometric hypotheses carrying a reliability estimate calibrated where scoring was on the experimental tier. That is what the crystal tier is for: it fixes what a given level of agreement is worth, so the corpus can be used as supervision rather than only as unlabelled geometry. A user whose requirement is volume of predicted geometry rather than measured separation might be better served by SAIR[8] or GatorAffinity-DB[10], which carry more systems but at the same time no experimental tier.

Two properties bound what the resource can support. The protein universe is inherited from a human protein–protein interaction gold standard[27], whose accessions were selected for sequence non-redundancy rather than for structural coverage. It contributes 906 accessions against 31,193 distinct ligands. Two accessions dissimilar in sequence can still present similar binding sites. The teacher configurations are throughput-oriented, chosen to reach corpus scale rather than to maximise per-system accuracy, so the coordinates record what those settings produce and not what the methods achieve when configured for a single target. Figure 1 shows the construction pipeline and Table 2 the per-arm settings, recovered from each run’s own configuration record.

**Fig. 1.**
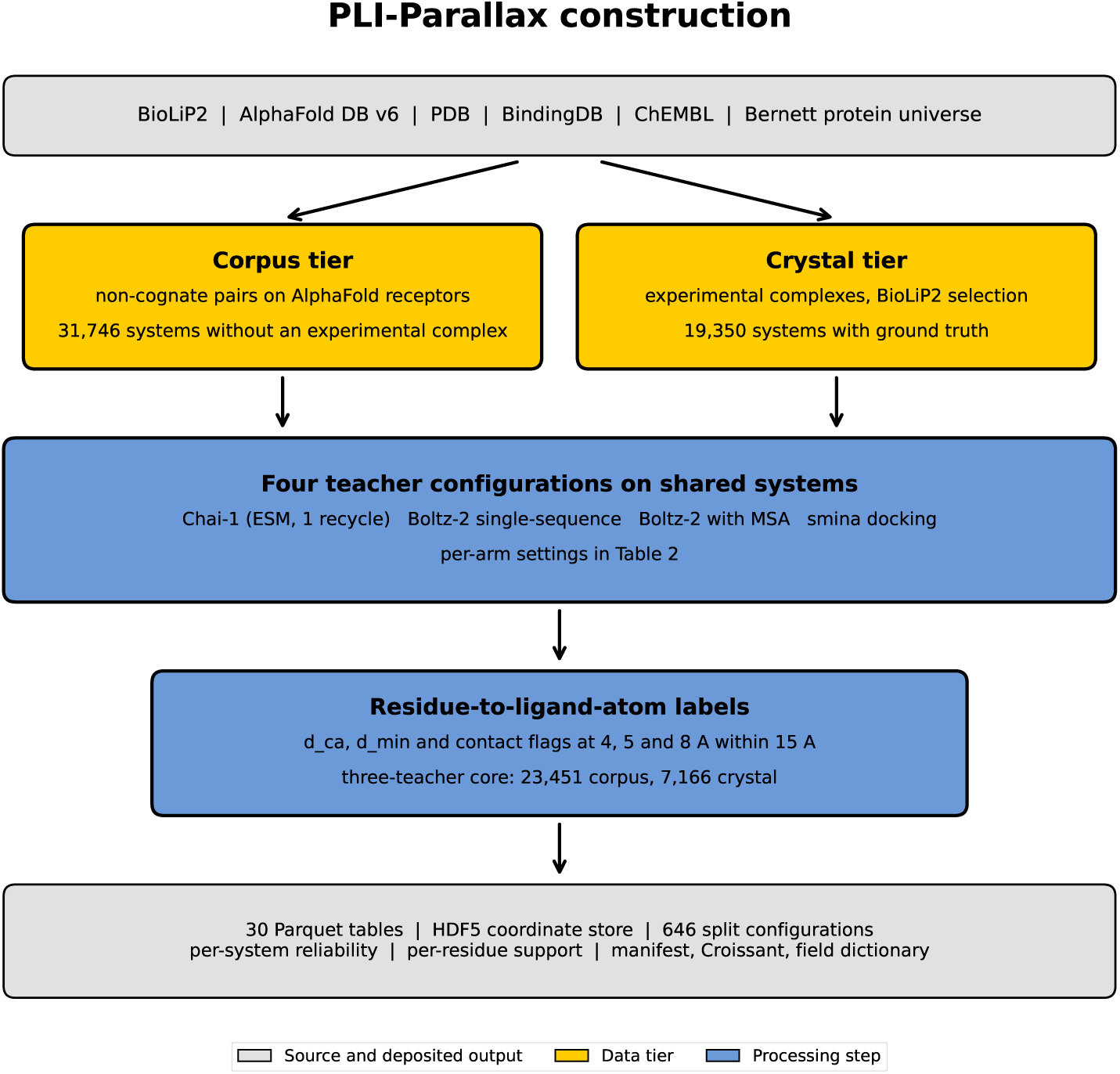
Construction of the deposit. Source databases supply the protein and ligand universe and the experimental coordinates. Systems divide into a corpus tier on AlphaFold receptors and a crystal tier from BioLiP2. Four teacher configurations run over shared systems, with per-arm settings given in Table 2. Labels record two distances per residue and ligand atom, of which only the minimum heavy-atom distance is subject to the 15 Å cutoff. The deposited output comprises the coordinate store, the distance tables, the split family, the reliability annotation, the per-residue support tables and the integrity metadata.

**Table 2.** Prediction arms. Each row is one run with its own configuration record, and settings were read from that record and not from documentation. Arms are not poolable across rows even where the model is the same, since alignment conditioning and recycling depth differ. The Systems column counts the deposited arm for the cofolding rows and attempted systems for the docking rows, which is the one arm that records attempts per system. 11,290 of 11,543 crystal-tier attempts produced a pose, and has contacts flags the difference per system.

| Arm | Model | Tier | Alignment | Recycles | Systems |
| --- | --- | --- | --- | --- | --- |
| 1 | Chai-1 | corpus | none, ESM | 1 | 23,485 |
| 2 | Chai-1 | crystal | none, ESM | 1 | 9,410 |
| 3 | Boltz-2 | corpus | none | 1 | 23,494 |
| 4 | Boltz-2 | crystal | none | 1 | 8,725 |
| 5 | Boltz-2 | corpus | ColabFold | 3 | 161 |
| 6 | Boltz-2 | crystal | ColabFold | 1 | 1,755 |
| 7 | smina | corpus | not applicable | – | 31,713 |
| 8 | smina | crystal | not applicable | – | 11,543 |

## Methods

### Overview

The construction has three stages, namely source selection, running the teacher arms, and deriving labels and annotations from what they produced. Coverage is not uniform across teachers in either tier, since each arm has its own failure modes, and the outcome is recorded per system in the deposit. Table 2 gives the per-arm settings and Figure 1 the pipeline.

### PLI-Parallax construction

#### Source Selection and Provenance

The crystal tier was defined by system selection from BioLiP2[25], which supplies, for each entry, a PDB identifier, a ligand chemical component identifier and a chain. Coordinates were read from the corresponding Protein Data Bank[26] entry rather than from BioLiP2 itself, so no BioLiP2 field value is redistributed here, the resource functioning as a selection list from which the labels derive through the pipeline described below. Table 3 lists every upstream source with its version or access date, its role in the construction, and its licence.

**Table 3.** Upstream sources. BioLiP2 supplies the crystal-tier system selection, and all crystal-tier coordinates are read from the PDB entry named by that selection.

| Source | Role | Version or date | Licence |
| --- | --- | --- | --- |
| BioLiP2 | crystal-tier system selection | 2026-06 | <sup>1</sup> |
| Protein Data Bank | crystal-tier coordinates | 2026-06 | CC0 |
| AlphaFold DB | corpus-tier receptors | v6, 2026-05 to 2026-08 | CC BY 4.0 |
| BindingDB | corpus-tier pair selection | 2026 | CC BY 4.0 |
| ChEMBL lineage | corpus activity threshold | (see Methods) | CC BY-SA 3.0<sup>2</sup> |
| Bernett et al.[27] | corpus protein universe | 2024 | CC BY 4.0<sup>3</sup> |

<sup>1</sup>The BioLiP2 source code is BSD-licensed while the data carries no explicit licence. It is used here as a selection list and no BioLiP2 field value is redistributed, so the coordinates in the deposit are governed by the PDB terms given below.
<sup>2</sup>Used as a selection threshold only. No ChEMBL-derived value is redistributed in the deposit, so the share-alike term does not propagate to it.
<sup>3</sup>The gold-standard split is distributed under CC BY 4.0 through figshare. Only the accession list is used, and no field value is redistributed.

Corpus receptors are AlphaFold Protein Structure Database[24] models at version 6, fragment F1, one model per UniProt accession, retrieved as AF-<accession>-F1-model v6.pdb. Corpus protein–ligand pairs were assembled from BindingDB[28], retrieved 2026, and from an upstream compilation of activity measurements. A pair entered the corpus where a pChEMBL value of at least 6.0 was recorded. The merge signature of the surviving source files is consistent with a two-feed assembly and identifies the activity values as ChEMBL-derived[29], but the specific release and retrieval route could not be recovered from the artifacts that survive on the processing host. No pChEMBL value is redistributed in the deposit, since the threshold determined which pairs appear rather than what is stored.

The dates in Table 3 are retrieval dates and not content coverage. A curated source is frozen before it is released, so the crystal-tier selection retrieved in 2026 carries entries deposited up to an earlier point, which Technical Validation states where it bears on the training-cutoff analysis.

The protein universe was inherited from the human protein–protein interaction gold standard of Bernett et al.[27], from which accessions with an available AlphaFold model were retained. That set was partitioned to control sequence-similarity leakage in interaction prediction and not for binding-site diversity, so the inheritance brings a non-redundant sequence universe and no guarantee of pocket coverage. Table 3 records what the corpus spans and Technical Validation reports the entity space it resolves to.

#### System Definition and Filtering

A system is a protein–ligand pair carrying a stable identifier. Crystal-tier identifiers take the form pdb ccd chain, while corpus-tier identifiers index an accession and a compound. Corpus construction is non-cognate by design, since a compound is modelled against a receptor with which it has never been co-crystallised, so the resulting labels record where a predictor places the ligand rather than where it is known to bind. Each teacher covers a different subset of each tier, for reasons recorded in Table 2.

The counts vary by denominator throughout this manuscript in a way that is not noise. The crystal three-teacher core is 7,166 systems by teacher coverage. Of those, 7,129 carry contact labels from all three arms and form the cohort on which contact accuracy is computed, while 7,134 carry an agreement value and appear in the reliability annotation. The corpus tier follows the same pattern, its three-teacher intersection standing at 23,451 systems of which 23,433 carry an agreement value. A system enters each denominator when the quantity that denominator defines can be computed for it, so the counts fall as the requirement tightens. Separately, the corpus spans 906 distinct accessions across all teacher arms, of which 678 carry cofolding coverage, resolving to 896 sequence clusters at 40% identity. These are measurements of different quantities and should not be substituted for one another.

The cofolding arms were run under a receptor length cap. Every accession they cover is at most 800 residues, while the 228 accessions they do not cover have a median of 1,101 residues and lie above the cap in 97.4% of cases. Those systems carry docking coordinates and no cofolding coordinates, which is the reason the docking arm is the largest on the corpus tier.

#### Applicability Domain

The three-teacher intersection is not a representative sample of the crystal tier, and users benchmarking on it should know in which direction it is biased. Core systems have a median of 414 residues against 1,344 for non-core systems, a median of 22 ligand heavy atoms against 44, and 5.5% metal content against 19.0%. Several ligand chemical component codes have a core fraction of exactly zero while appearing in the full tier, and Table 4 lists a selection of the larger ones.

**Table 4.** Composition of the three-teacher core against the rest of the crystal tier. Ligand classes listed below the rule appear in the crystal tier and are absent from the core entirely, and counts are systems. The list is a selection and not the complete set of such classes. A model selected on the core and applied to large or metal-bearing systems is extrapolating.

| Property | Three-teacher core | Non-core |
| --- | --- | --- |
| Residues (median) | 414 | 1,344 |
| Ligand heavy atoms (median) | 22 | 44 |
| Chains (median) | 2 | 4 |
| Metal-containing (%) | 5.5 | 19.0 |
| Interface binder (%) | 14.6 | 32.4 |
| *Ligand classes with zero core fraction* |  |  |
| SF4 | 0 | 535 |
| CLA | 0 | 375 |
| CO | 0 | 142 |
| NAI | 0 | 87 |
| DD6 | 0 | 84 |
| CHL | 0 | 74 |
| BCL | 0 | 46 |
| CU1 | 0 | 43 |
| B12 | 0 | 43 |
| FDA | 0 | 30 |

One row of Table 4 matters more than the others for a resource of distance labels. The core contains an interface binder in 14.6% of systems against 32.4% outside it, a depletion of more than twofold. Counting contacted chains rather than interface status points the same way, with 49.0% of core systems contacting exactly one chain against 12.3% outside. A contact model selected on the core is selected against multi-chain binding sites.

#### Cofolding Arms

Chai-1[19] (version 0.6.1) was run with one trunk recycling step, 80 diffusion timesteps and ESM embeddings enabled, producing five diffusion samples per system, of which the rank-0 sample supplied the labels. All five are deposited. Boltz-2[20] (version 2.2.1) was run in several configurations that differ in whether a multiple sequence alignment was supplied and in the number of recycling steps. Alignments, where used, were retrieved from the ColabFold MMseqs2 server[30] with greedy pairing.

Recycling depth was not uniform across arms, and an earlier release note for this deposit stated it incorrectly for one of them. Reported for every arm is the value recorded in that run’s own predict args block rather than the value intended at launch. The two crystal arms named single-sequence and alignment-conditioned in Table 2 both ran one recycling step, so the comparison between them varies only whether an alignment was supplied. The alignment-conditioned corpus pilot ran three recycling steps, and the arms not deposited here, run as controls during construction, ran three. Recycling does not apply to the docking arm.

The reliability annotation reads agreement among three arms of deliberately unequal strength. The single-sequence configuration is the weakest of them by every accuracy measure reported here, and it is retained for that reason: agreement between arms that differ in accuracy separates easy systems from hard ones, where agreement between arms of similar strength would not. The alternative without it would be a consensus over two configurations of one lineage and one docker, on a different intersection, which is a narrower signal and not the same systems.

Several counts describe the deposited alignment-conditioned arm and they measure different things. 1,755 systems produced a pose and have a metadata row. 1,751 of those carry contact rows in the label table, which is the denominator of its contact-accuracy figure. 1,743 were scored for physical validity, and 456 resolved to a symmetry-corrected root-mean-square deviation, which requires an atom-graph isomorphism to the reference ligand and fails where none exists. This arm is not one of the three whose mutual agreement defines the core cohort, so its contact accuracy is computed on its own systems and is not comparable to the other three.

Cofolder predictions are conditioned on the protein the model itself predicted, not on any deposited receptor, so each arm carries its own receptor conformation and its own coordinate frame. Comparing a pose across teachers therefore requires superposing on shared residues first.

Table 2 gives the arms, their conditioning and their coverage. The teacher set is a snapshot of what was available when the arms were generated, not a selection of the best models at time of publication. Open alternatives released since, among them Protenix[31], are not represented here. Extending the teacher axis is deferred to a subsequent version of the deposit rather than applied retroactively, since adding an arm changes the intersection on which every consensus quantity is computed.

#### Docking Arm

The docking arm used smina[22], a fork of AutoDock Vina maintained separately from the AutoDock project. Smina carries no semantic release number, so the arm is pinned by its base version, Vina 1.1.2, together with the build date 15 October 2019. Receptor and ligand were converted to PDBQT with Open Babel.

Ligands were built from the chemical component SMILES with RDKit 2026.03.3. A single conformer was embedded at a fixed random seed, falling back to random coordinates where embedding failed, and minimised with MMFF. No protonation adjustment or tautomer enumeration was applied, so each ligand carries the protonation state and tautomer its chemical component SMILES specifies, and RDKit defaults govern throughout. Since hydrogens are excluded from the labels on both sides, that choice reaches the distance tables only through the heavy-atom count of the ligand. The docked ligand’s internal geometry therefore originates from RDKit rather than from any experimental structure, which is the correct procedure for docking and means its bond lengths and ring geometry satisfy a validity check by construction.

Receptors retain every heteroatom residue that AutoDock can type. Residues carrying an element outside the typeable set were removed whole rather than atom-wise. Removing only the untypeable atom would leave an incomplete residue in the pocket, and atom typing is inferred from geometry, so the fragment would be typed as though it were complete. The filter removed residues from 167 systems. Of those residues, 126 across 95 systems fall inside the search box, and the remainder lie outside it where they cannot influence the result.

Docking used a 22 Å cubic box and a fixed seed, at exhaustiveness 8 with three output modes on the corpus tier and exhaustiveness 4 with one output mode on the crystal tier. On the crystal tier the box was centred on the deposited ligand centroid, one box per system. On the corpus tier no experimental pose exists, so the candidate pockets from the ensemble annotation were ranked by the rank score recorded with them and the highest-ranked five docked independently, the pose with the lowest predicted affinity retained and the originating pocket recorded in pocket rank. That choice is not a formality. The top-ranked pocket wins on 49.9% of corpus systems and the fifth-ranked on 6.9%. The pocket ensemble covers the corpus tier only, so no multi-pocket search was performed on the crystal tier.

The crystal arm ran under a 120 second per-system cap. Of 11,543 attempts, 11,290 produced a pose, and the difference is dominated by systems that reached the cap, biased toward the largest since a larger search space takes longer to exhaust. That bias runs in the same direction as the small-receptor composition of the three-teacher core and should be read alongside it. Symmetry-corrected deviations were computed both for the whole posed arm and for the subset that the superseded build also scored, and the deposited scoring output carries both.

The consequence should be stated. On the crystal tier the docking arm receives the site and the cofolding arms do not, so the three arms are not exchangeable observers there. On the corpus tier none receives it. That asymmetry is a difference between the tiers beyond the non-cognate construction, and it bears on any quantity computed from agreement among the three.

The docking arm deposited here was regenerated from an earlier build, and the reason bears on how the arm should be read. The receptor preparation routine returns early when its output file already exists, which is correct in the context it was written for, where the output path is keyed by accession. The calling script passed a single output path per shard while supplying a different receptor at each iteration, so every system after the first in a shard was docked against the first system’s protein. The corpus arm is keyed by accession and was not affected. On the crystal tier the defect reached 11,529 of 11,543 systems, or 99.88%, and that build is not deposited. The arm described here was re-docked from scratch with a per-system receptor path. Median symmetry-corrected ligand root-mean-square deviation falls from 8.80 to 4.15 Å on the 6,485 systems scored in both, and the median docking score falls from *−*0.0009 to *−*7.64 kcal mol*^−^*^1^, the earlier value reflecting poses that were not scoring a pocket at all. The identifiers of the affected systems are deposited alongside the arm.

#### Label Definition

The distance tables are derived from the coordinate store at a fixed cutoff and a fixed pair definition, and both quantities they record are recomputable from it. For each system, every ligand heavy atom is compared with every protein heavy atom. A residue enters the table where the minimum distance from any of its heavy atoms to any ligand heavy atom is at most 15.0 Å. For each qualifying residue and ligand atom the table records d min, that minimum heavy-atom distance, and d ca, the distance from the ligand atom to the residue C*α*. Because the cutoff applies to d min, d ca may exceed 15.0 Å, and filtering on it would silently discard rows. Where a residue carries no C*α*, the first heavy atom substitutes, so d ca is a C*α* distance for all but those residues, which are present in the tables rather than dropped. Hydrogens are excluded on both sides. Boolean contact flags are derived at 4, 5 and 8 Å on d min.

Distances are stored at half precision, so values lie on a grid whose spacing doubles at each power of two: 0.0039 Å just above 4 Å, where the contact flag sits, and 0.008 Å near 12 Å. An analysis whose outcome turns on a threshold should allow one step of margin at that threshold.

Residue indices are zero-based within the receptor model and join to the residue metadata tables. Ligand atom indices follow the SMILES reference ordering shared by the prediction arms, while the ground-truth arm follows deposited file order, in which unmodelled atoms are absent, so the two orderings are not interchangeable.

The rdkit parse ok field carries a name narrower than its use. It records one condition, that the SMILES parsed, and parsing is the first of several a system must meet before it can be folded. Read as a parsing outcome the field is correct. Read as a foldability flag it is permissive, and the 50.5% it reports on the ground-truth arm is the parse rate rather than the fold rate.

#### Split Construction

646 train, validation and test configurations are provided across seven families and both tiers. A configuration is identified by four things: the family, which entity axis is held out; the protein identity threshold at which sequences were clustered; the ligand clustering variant; and the seed, an integer from 0 to 4. The families follow the C1, C2 and C3 distinction of Park and Marcotte[32], subdivided here, and five of the seven vary the entity axis while two vary a different one. C1 holds out neither, so both the protein and the ligand cluster are seen in training, while C2p holds out the protein and C2l the ligand. C3 and C3comp hold out both, the first by block assignment and the second by connected component. Two further families vary a different axis, T1 partitioning by deposition date and LO holding out lead-optimisation series. The family label states which entity a claim generalises over, so a configuration is chosen against the claim being made rather than defaulted.

The five entity-axis families are provided at three protein identity thresholds on the corpus tier and two on the crystal tier, giving 375 corpus and 250 crystal configurations, with 6 temporal and 15 lead-optimisation. Every corpus configuration assigns all 31,746 systems. The crystal configurations assign 18,458 of 19,350, leaving 892 unassigned, because their ligand clusters are large enough that placing one in a single fold would unbalance every configuration built from it. 748 of those carry haem and five ligand codes account for most of the rest, the same metal macrocycles and cofactors Table 4 records as under-represented in the three-teacher core. One cause bears on both.

Protein clusters were computed with MMseqs2[33] (commit 17b688d2) using easy-cluster at coverage 0.8 and cluster mode 2, at three sequence identity thresholds. Clustering keys differ by tier. The corpus tier clusters on UniProt accession, one receptor per accession, while the crystal tier clusters on the sequence hash of the key chain, since 14,852 distinct key-chain sequences occur across 19,350 systems and an accession key would merge sequences that differ. At 30% and 50% identity the crystal tier resolves to 7,149 and 10,714 clusters.

Ligand clusters were computed by Butina leader clustering over 2048-bit Morgan fingerprints of radius 2, at four Tanimoto cutoffs, with a Bemis–Murcko[34] generic scaffold variant provided alongside. The fingerprint variant at cutoff 0.40 is the one quoted for prospective claims, the scaffold variant is more permissive, and the two partitions are not nested. Crystal-tier ligands were resolved from the chemical component dictionary SMILES, of which 952 are acyclic and therefore carry no Murcko scaffold. The two tiers treat these differently. On the crystal tier they were assigned singleton clusters, so that a scaffold-based family does not group unrelated chemistry. On the corpus tier they follow the conventional scaffold split and share one cluster, so a user selecting a scaffold-variant corpus configuration holds out that cluster as a unit. The policy is recorded per variant in the deposited clustering report. Seeds are the integers 0 to 4 for every family and threshold combination, so a configuration is fully identified by family, threshold, ligand variant and seed.

Seen and unseen are evaluated at cluster level throughout rather than at exact entity level. 98.0% of corpus ligands occur in exactly one system, so an exact-identity condition would cap the warm classes at a few hundred systems and the family axis would collapse. Cluster crossing is zero on every axis a family holds out and deliberate on every axis it does not, which is what warm split refers to. What that delivers differs by axis and by tier. On the corpus tier a warm protein axis returns the same accession, with an exact-seen fraction of 1.000, while a warm ligand axis returns a cluster neighbour rather than the same compound, at 0.05. The crystal tier reverses it, at 0.50 and 0.90. A key-chain sequence cluster holds several distinct sequences, while a recurring cofactor appears in hundreds of systems, so the two axes behave oppositely there. On any held-out axis the exact-seen fraction is 0.000. Reporting these fractions rather than asserting a threshold is deliberate, since a threshold claim is checkable only against the construction that made it while a measured fraction is checkable against the deposit.

Cross-fold sequence identity is reported under a coverage-normalised definition, the alignment identity scaled by alignment length over the shorter sequence. The raw local identity reported by the aligner must not be read as a leakage figure, because a short perfect local match is reported as complete identity regardless of coverage. Technical Validation reports the measured values under both definitions and per family, since the figure is interpretable only where the protein axis is held out.

The gap between the two definitions is large enough to change a conclusion. Reporting the raw figure would describe the splits as heavily contaminated when the contamination is an artifact of short high-scoring local alignments, and Technical Validation gives the measured gap. The splits are therefore described as redundancy-reduced at a stated maximum rather than as leakage-controlled, following the reporting convention established for protein–ligand data by PLINDER[6], LP-PDBbind[5] and CleanSplit[35]. The weaker word is deliberate. A leakage-controlled claim asserts that a threshold was met and cannot be checked without rebuilding the partition, while a maximum reported per configuration is a value in a deposited table that a reader can read off.

#### Coordinate Store

Predicted and experimental coordinates are provided in an HDF5 store, one group per system. Coordinates are mean-centred per system, with the centroid retained at single precision, and quantised to signed 16-bit integers at a scale of 100 units per Å, giving a resolution of 0.01 Å. The round trip was verified at 0.002 Å per coordinate, so a distance computed from two of them carries about twice that. Mean-centring is required rather than conventional, since absolute coordinates would exceed the representable range at this scale.

The store does not cover every system. It holds 31,617 of the 31,746 corpus systems and 17,368 of the 19,350 crystal systems. On the corpus tier all 129 omissions failed to anchor a reference ligand. On the crystal tier 1,720 failed to anchor, 260 carried no usable reference at all, and two raised a parsing exception on deuterium-substituted coordinates. The skipped systems retain their distance-table rows, so a user recomputing distances from the store recovers a subset of the tables rather than all of them.

The representable range at this scale is 327.67 Å either side of the centroid, which 89 systems reach, on the crystal tier through assembly size and on the corpus tier through extended low-confidence tails in the fallback receptor. Shell atoms and every label-referenced residue lie inside the range in all of them, so no deposited label is affected, and the affected identifiers ship as metadata/int16 saturated systems.json.

Each system stores its ligand poses, up to 5 per teacher, on a fixed teacher axis with experimental ground truth at index zero. A score per pose is stored alongside, with a field recording its units and direction, since the teachers report different quantities. For each teacher the store holds the receptor C*α* trace for the whole chain and every heavy atom of a standard residue within 15.0 Å of the ligand, with element and residue index. The residue enumeration admits only the twenty standard amino acids, so a metal ion, a cofactor or a modified residue inside the cutoff is absent from the shell even where it contacts the ligand. Receptor coordinates are stored per teacher because each cofolding model predicts its own receptor conformation.

Two further points are stored per shell residue. A side-chain centroid, the geometric centre of the residue’s heavy atoms excluding backbone N, C*α*, C and O, fixes the direction and extent of the side chain, which the C*α* trace alone does not express. A single bead at the side-chain centre is the representation from which all-atom geometry has been reported to recover most accurately[36], and reconstruction from reduced representations is long established[37]. Glycine carries no side-chain heavy atom, and its centroid is taken over the four backbone heavy atoms instead of being placed at the C*α*. That departs from the convention used in that work, where a coincident point is harmless, because a coordinate coincident with a stored C*α* produces a degenerate zero-length pair for a consumer computing distances. The second point is the backbone carbonyl carbon, which fixes the direction in which the chain leaves the residue and is present in every residue including chain termini, so it requires no mask.

The shell radius matches the label cutoff, so the store contains every atom that could contribute to a label. A residue entering the label table may nonetheless carry individual atoms beyond the shell.

#### Reliability Annotation

Each system carries a predicted label accuracy with an interval, provided as a released field rather than as an analysis of the labels. Inter-teacher agreement was computed as the mean pairwise Jaccard index between the 4 Å contact residue sets of Chai-1, Boltz-2 single-sequence and the regenerated docking arm, so no deposited agreement value derives from the superseded build. The three pairs do not contribute equally and the deposited tables show how they differ. On the corpus tier, where no arm receives the site, Chai-1 agrees with the single-sequence arm at 0.1224 and with the docking arm at 0.1218, which are the same figure, while the pair involving the single-sequence arm and the docker reaches 0.0798. Two models that share the AlphaFold3 design and a training archive therefore agree no more with each other than either does with a physics-based docker, so the signal is not an artifact of shared lineage. On the crystal tier the ordering changes. Chai-1 against the docking arm rises to 0.3362 while the two cofolders reach 0.1909, and the arm whose agreement rises most is the one given the site there. That asymmetry is a property of the crystal tier and not of the teachers, and it is the reason the agreement measured there is between observers of unequal input.

Where accuracy against ground truth is measurable, systems were split in half at a fixed seed. Accuracy is the same Jaccard overlap used for agreement, computed between an arm’s contact residue set and the ground-truth set rather than between two arms, so both axes of the regression are the same measure. An isotonic regression[38] of accuracy on agreement was fitted on one half, and the 90th percentile of absolute residuals on the held-out half gives a split-conformal[39] half-width. Empirical coverage equals nominal at both levels reported by construction, since the half-width is the corresponding quantile of those residuals. The fitted mapping was then applied to the corpus tier. Two error figures are reported. Held-out error is given as a five-fold cross-validated mean absolute error so that it does not depend on which half the fit received, and the deposited calibration split gives 0.0805 against 0.1752, within 0.002 of the cross-validated pair. Conformal prediction has been used in this domain as an alternative to applicability-domain determination[40], which is the role it plays here. The interval is a marginal split-conformal bound calibrated on the crystal tier.

Crystal-to-corpus is a genuine covariate shift, so the guarantee does not transfer unweighted, and the reweighting that would recover it is not applied because the estimate it requires is unreliable where corpus systems concentrate. What transfers is the ordering and not the bound, so the field ranks corpus systems by expected label quality rather than bounding any individual system’s accuracy.

#### Software

Structure prediction used Boltz-2[20] 2.2.1 and Chai-1[19] 0.6.1 under PyTorch 2.12.0 on Python 3.12.13, on single consumer GPUs over several months at fixed seeds recorded per arm. Docking used smina[22] with Open Babel[41] 3.1.1. Cheminformatics used RDKit[42] 2026.03.3 and structure parsing gemmi[43] 0.6.5. Tabular storage used pyarrow 25.0.1 and the coordinate store h5py 3.16.0. Clustering used MMseqs2[33], physical validity PoseBusters[44] 0.6.5, symmetry-corrected deviations spyrmsd[45] 0.9.0, and the reliability calibration scikit-learn[46] 1.6.1. Alignments were retrieved from the ColabFold[30] MMseqs2 server.

Four of these could not be pinned to a release number. Smina carries none and is identified by its build date, 15 October 2019, on an AutoDock Vina[47] 1.1.2 base, as is Open Babel by its own build date 31 March 2024 alongside the release above. MMseqs2 reports a commit rather than a release and is identified here as commit 17b688d2. The ColabFold server exposes no client-queryable version and is identified by endpoint and retrieval date. The Chai-1 and spyrmsd versions are the values their runs recorded. They are reported as recorded rather than re-verified, since the relevant version is the one that produced the coordinates and not the one a fresh installation would supply.

#### Data Records

The dataset is deposited at Zenodo under DOI 10.5281/zenodo.21560088[48], released under CC BY 4.0. The code that produced it is at https://github.com/ThorKlm/PLI-parallax under Apache 2.0 and archived at Zenodo under 10.5281/zenodo.22258211. The deposit is 6.16 GB, of which the tabular layer is 1.14 GB and the coordinate store 5.01 GB.

The deposit is layered, and the layers are not alternatives. A coordinate store holds the geometry and is the primary artifact. Distance tables hold a materialised view of that store at a fixed cutoff and a fixed pair definition, provided so that a user wanting distance labels need not compute them. Derived annotations key into both through the identifiers described below: the split family, the pocket annotation, the per-residue support tables, and the reliability estimate that the two tiers exist to calibrate. The derivation is checked rather than assumed: recomputing both distance columns from the store reproduces the deposited values to the quantisation floor across the full store, while nothing in the store is recoverable from the tables. The store does not cover every system, omitting 129 on the corpus tier and 1,982 on the crystal tier for the reasons Methods gives, so a user recomputing from it recovers a subset of the tables rather than all of them.

#### Coordinate store

The store is a set of HDF5 files, sharded for parallel construction and otherwise independent, with one group per system at the root. Coordinates are mean-centred per system and stored as signed 16-bit integers at a scale of 100, giving 0.01 Å resolution. The centroid that was subtracted is retained per system as centroid xyz in single precision, so absolute positions are recoverable.

Each system group carries a fixed teacher axis of length five, ordered as experimental ground truth, Chai-1, Boltz-2 single-sequence, Boltz-2 alignment-conditioned, and smina. The axis is fixed so that a consumer indexes by position rather than by name, and a system that lacks a teacher carries a false entry in ligand valid rather than a missing dataset. Which teachers a system actually carries is recorded per system rather than inferred from the file-level axis. Ligand coordinates are held as ligand coords, shaped by teacher, pose, atom and axis. Only Chai-1 uses more than one pose slot, storing all 5 diffusion samples. Every other teacher stores one. Index zero is the sample the distance labels were derived from, since recomputing d ca from it returns the deposited value to the quantisation floor and recomputing from any other index does not. The remaining four samples are retained for users who want the spread and carry no deposited score. pose score is populated for the docking arm alone, with score kind recording its units and direction. The cofolding arms declare no score, since a cofolder confidence and a minimised affinity are neither comparable nor ordered the same way.

Receptor geometry is stored per teacher, because each cofolder predicts its own protein and the conformations are not interchangeable. Under protein/<teacher>/ the store holds ca, the C-alpha trace of the full receptor, and shell coords, every heavy atom of a standard residue within 15 Å of the ligand, with shell elem and shell res index giving element and residue assignment. The qualification is loadbearing. The residue enumeration admits only the twenty standard amino acids, so the shell contains carbon, nitrogen, oxygen and sulfur and nothing else, and a metal ion or a cofactor inside the cutoff is absent from it even where it contacts the ligand. Two further arrays are held per shell residue, sc centroid with its sc valid mask and pep c, both defined in Methods.

The frames are not shared. Comparing the crystal and single-sequence receptors of the same system residue by residue gives a median C*α* offset of 50.4 Å, so a comparison across teachers requires superposition first.

The three per-residue points are stored as separate datasets and are never merged. A consumer wanting a C-alpha trace alone, or C-alpha with a side-chain bead, selects columns instead of converting formats. Table 5 lists the reduced views this supports and the derivation of each.

**Table 5.** Reduced views of the coordinate store. Each is a column selection or an index subset and not a format conversion, so a consumer reads only what it needs. Distances are derived at read time and none is stored. The distance tables are the exception, and they are themselves the ligand-to-residue view materialised at a fixed cutoff.

| View | Derivation |
| --- | --- |
| C-alpha trace, full receptor | `ca` |
| C-alpha trace, pocket | `ca` subset by `shell_res_index` |
| Two beads, pocket | the above with `sc_centroid` |
| Three beads, pocket | the above with `pep_c` |
| All heavy atoms, pocket | `shell_coords` with `shell_elem` |
| Ligand heavy atoms | `ligand_coords` by teacher and pose |
| Residue-residue distances | pairwise over any protein view |
| Ligand-ligand distances | pairwise over `ligand_coords` |
| Ligand-to-residue `d_ca` | cross-pairwise, `ca` against ligand |
| Ligand-to-residue `d_min` | minimum over `shell_coords` grouped by residue |

The shell is defined by distance to the ligand, so a residue that contributes a contact may have individual atoms outside it. The store contains every atom that can produce the minimum recorded in the distance tables. It does not contain every atom of every contacting residue.

#### Distance tables and metadata

Distance tables are Parquet, one file per arm and tier, with a uniform eight-column schema. Each row is a residue and ligand-atom pair within 15 Å, recording the two distances defined in Methods, d ca and d min, and boolean flags at 4, 5 and 8 Å derived from the second. Rows are addressed by system id, res row and atom idx. Recomputing both distances from the store reproduces the deposited values to 0.0040 Å, which is the quantisation floor of the store rather than a disagreement.

Metadata tables share a uniform 21-column schema across all teachers and both tiers, giving 129,636 rows in total. Not every column is populated for every teacher: conf1 and conf2 carry values on the Boltz-2 corpus arm alone and are null on the eight other files, affinity and pocket rank are null on the cofolding arms, and several structural columns are populated on the crystal tier alone. The schema is uniform. The content is not. A metadata table carries one row per attempted system while its distance table carries rows only where a pose was produced, so the two counts differ where a teacher failed on some systems. The docking arm on the crystal tier is the largest case: 11,543 attempted rows against 11,290 systems with contacts, the difference flagged by has contacts.

Two identifier conventions differ between tables and both are load-bearing. The metadata tables key proteins as protein id while the residue table keys them as accession, and although the values are the same UniProt[49] accessions and the join is direct, the column names are not interchangeable.

Table 6 lists every deposited file with its format, row count and size. The field dictionary is not reproduced here. It ships with the deposit as FIELDS.md and FIELDS.json, covering 235 fields across 14 record sets with the type, unit and definition of every column, and it is generated from the same file as the Croissant description rather than written by hand.

**Table 6.**
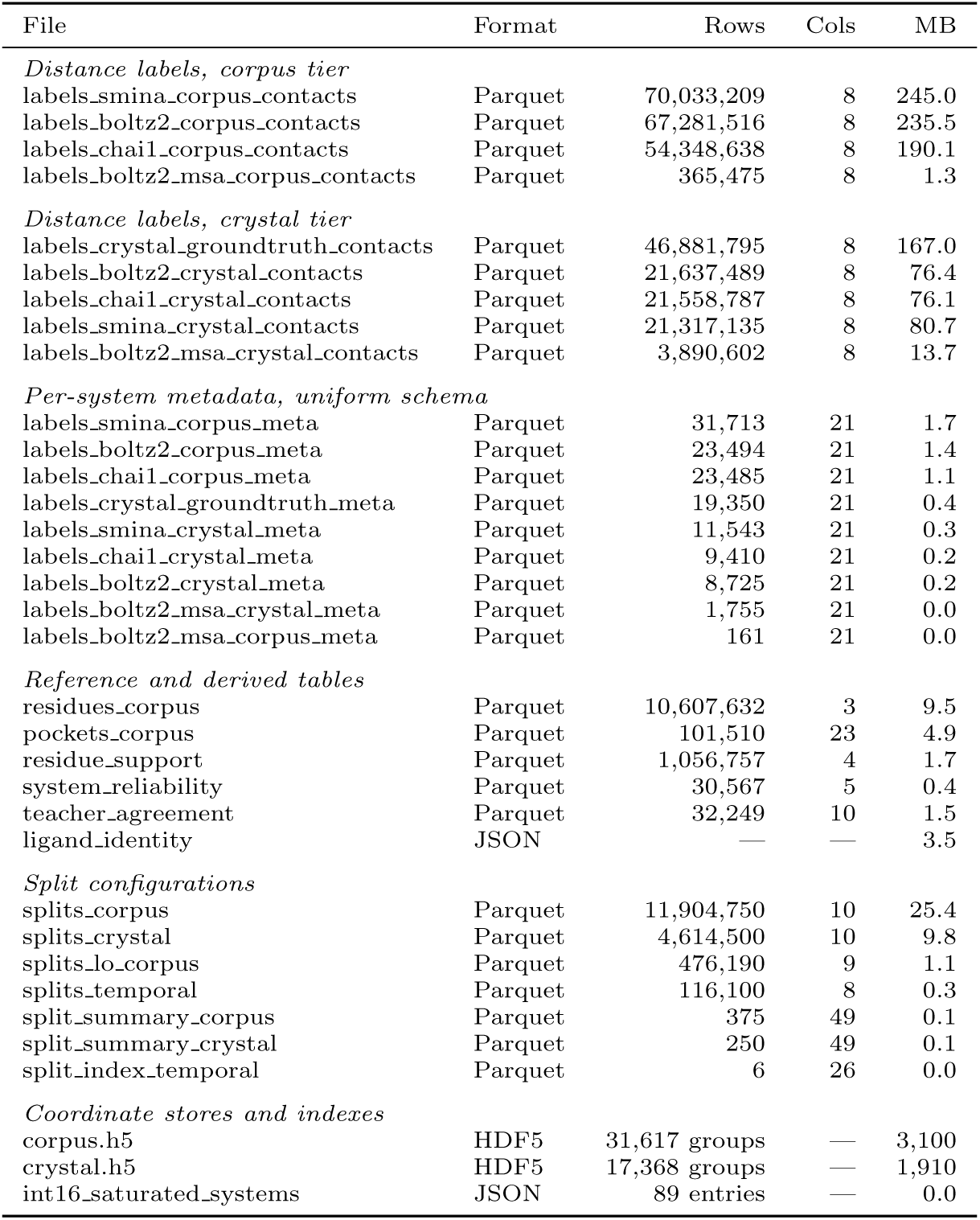
Deposited files with their format, row count, column count and size as archived. Documentation and integrity metadata are described in the text and not listed here. Contact rows total 307,314,646 and metadata rows 129,636, which together with the reference, derived and split tables give 336,385,168 rows across the tabular layer.

#### Derived annotations

The split family ships as seven Parquet files, one fold table and one summary table per tier together with the temporal and lead-optimisation tables, keyed by split tag and system id. Across both tiers they carry 646 configurations, constructed as Methods describes. The fold table gives one row per system and configuration. Three summary tables give one row per configuration with its diagnostics, covering the 375 corpus and 250 crystal entity-axis configurations with classifier separation scores and the 6 temporal configurations with exact-entity overlap fractions. The 15 lead-optimisation configurations ship as a fold table alone and carry no summary.

The reliability annotation gives one row per system, recording a predicted label accuracy and a split-conformal interval at the stated coverage. It is calibrated on the crystal tier, where accuracy is measurable, and applied to the corpus tier, where it is not. Methods states what that transfer does and does not license.

The pocket annotation gives 101,510 pockets over 20,404 accessions, capped at five per receptor and averaging 4.98, so the cap binds on almost every receptor. Each pocket carries a centroid and 18 features including per-detector source flags, which with the three centroid coordinates and the identifier columns give the column count in Table 6. The three detectors are fpocket[50], which is geometric, and P2Rank[51] and VN-EGNN[52], which are learned, so a pocket found by more than one is supported by more than one kind of evidence. A combined rank score orders the pockets of a receptor and is the field the docking arm selected on.

Two limits apply. The annotation covers the corpus tier alone, since it was computed on the AlphaFold receptors, and the crystal tier has no equivalent. And the step that merged the three detectors did not retain per-pocket residue membership, so membership is not deposited and the merge is one of the stages Code Availability records as not reproducible from the repository. Membership is derivable from the centroid at a radius of the user’s choosing, and the residue count recorded for the P2Rank-detected subset gives a reference for that choice.

The residue table maps each receptor to its sequence, one row per accession and residue index. It covers 20,404 accessions, which is the full AlphaFold receptor collection from which the corpus was drawn rather than the 906 that carry a system, so a row count divided by the corpus protein count is not a residues-per-protein figure.

Per-residue support tables record, for each residue in the three-teacher core, how many teachers place a ligand atom in contact with it. The fields are system id, tier, res row and n teachers asserting, the last an integer from one to three. Precision against ground truth as a function of that count is reported in Technical Validation. A separate table records geometric disagreement rather than contact-set overlap, one row for each of 32,249 systems carrying more than one teacher. Its fields are system id, tier, n teachers, the median and maximum pairwise ligand centroid offset, the median and maximum pairwise ligand root-mean-square deviation, the pocket residue count and its union across teachers, which are the basis on which the superposition was computed, and a note column recording why a value is absent where it is. Each teacher places its ligand against the receptor it predicted, so an offset computed without first superposing on the shared pocket residues measures the difference between two coordinate frames and not the disagreement between two poses. The deposited values are superposed, and a user recomputing them from the store must superpose as well.

#### Integrity

A SHA-256 manifest covers every deposited file, and an inventory records the row count, byte size and compression codec of each tabular artifact. Croissant metadata describes the tabular layer in machine-readable form. The manifest, the inventory, the Croissant description, the deposited FIELDS.md and the README schema section are all generated from one field definition file carrying 235 entries across 14 record sets, so the five cannot diverge. Generation order is enforced so that no artifact records a hash of a file a later stage rewrites. Croissant describes the 34 data files and not the six documentation files, which the manifest covers and which carry no schema to declare. A reader who extracts the archive can therefore verify every file against the manifest and every column against the dictionary without consulting this article.

#### Technical Validation

The checks below fall into two groups. The first establishes that the deposit is internally consistent and that the artifacts stand in the relationship Data Records claims. The second establishes what the labels are worth, and uses the crystal tier throughout, because that is where accuracy is measurable.

The corpus tier is not validated in the same way. No experimental complex exists for any of its 31,746 systems, so no corpus label can be scored against a measurement.

Reported instead is a reliability estimate calibrated where accuracy is measurable and transferred to where it is not. That is how prediction-derived resources report on their unmeasurable portion, the AlphaFold DB shipping a per-residue confidence in place of an accuracy it cannot measure[24, 53].

#### Integrity

Every deposited table satisfies its stated invariants to the precision at which they are stored. d min exceeds d ca on 195 of 307,314,646 rows, all in the ground-truth table and all by at most 0.0078 Å. In exact arithmetic the two are equal wherever the nearest heavy atom of a residue is its own C*α*; at half precision they round to adjacent grid points, and the excess is bounded by one step of that grid. Joining each distance table to its metadata table on system id resolves every row, and the uniform 21-column metadata schema concatenates across teachers and tiers without type coercion.

Recomputing d ca and d min from the coordinate store reproduces the deposited values to a median 0.0040 Å over 1,754 comparisons spanning the full store. The median rather than the mean is the right statistic here: on a ligand with symmetry-equivalent atoms the store resolves the match against a per-system anchor while the label extractor took the first substructure match, so the two disagree legitimately and the mean rises to 0.4352 Å. The median is the quantisation floor of the store rather than a disagreement between artifacts, and it is the evidence for the derivation relationship stated in Data Records.

The store’s own construction is verified per system rather than in aggregate. Chain identity is resolved by mmCIF chain name and never by position, each chain is required to match the recorded sequence and residue count exactly, and the residue remap is required to be a permutation of the full residue range. The deposited chain schema records the outcome of all three for every crystal-tier system and reports them satisfied on all 19,350.

The validation suite in the code repository runs 57 checks against the archived record and takes the deposit root as an argument. Fifty read only the deposited artifacts, and seven compare the coordinate store against the source mmCIF and AlphaFold files, reporting as skipped where those are absent. verify quoted figures v2.py recomputes 130 of the figures quoted in this article directly from the deposited Parquet and names the thirteen it does not reach. A reader can re-run both against the archived record instead of taking the numbers here on assertion.

#### Composition and applicability

The entity space is wider than the system count alone would suggest. 906 distinct accessions resolve to 896 sequence clusters at 40% identity, and the threshold is not load-bearing, since 30% and 50% give 886 and 900 clusters respectively. Ligand clustering runs over the 31,063 ligands carrying a resolvable structure, which is fewer than the 31,193 distinct ligands the corpus spans. Those resolve to 15,173 Bemis-Murcko generic scaffolds, of which 70.7% are singletons, and to 7,933 clusters under the fingerprint variant at Tanimoto 0.40 that the prospective families use. The largest scaffold cluster holds 274 members. The corpus is therefore a wide entity set rather than a narrow one inflated by repetition.

The applicability bias stated in Methods is recorded per system rather than only in aggregate. A user benchmarking on the three-teacher core can therefore identify which systems the core omits and filter accordingly, selecting on the metal and chain-count fields of the metadata schema for the two covariates that separate the cohorts most sharply.

#### Split leakage

All 646 split configurations were checked for cluster crossing on both entity axes. Every configuration crosses zero clusters on the axes its family holds out, and the deposited summary tables record the crossing and the exact-entity overlap per configuration, so the property is checkable rather than asserted. A warm axis crosses by construction, which is the definition of warm and not a leak.

Cluster separation is a construction property and says nothing about whether the resulting folds differ. Each configuration therefore carries a two-sample classifier score[54], the held-out accuracy of a discriminator trained to tell training entities from test entities, together with the score the same discriminator reaches on protein length alone. The second is the length-only baseline the first has to be read against, since a configuration whose protein-axis score sits at it has not made its proteins distinguishable by anything beyond size. The scores are reported as evidence that the shipped configurations separate as their family labels claim, and not as a measurement of any model.

On the ligand axis the corpus tier rises with family strictness overall, from 0.508 where both axes are warm to between 0.824 and 0.899 where both are cold. It does not rise in family order. Holding out the protein separates the ligand distributions more than holding out the ligand does, because a corpus in which 98.0% of ligands occur in exactly one system has little ligand structure to remove. The crystal tier inverts entirely, reaching 0.962 on the cold-ligand family against 0.618 for the same family on the corpus, since holding out a cofactor that recurs across hundreds of crystal-tier systems removes its proteins with it. On the protein axis the corpus sits at its own length baseline in every family, so partitioning proteins does not by itself make them distinguishable, while the crystal tier reaches 0.7798 against a baseline of 0.5735. Figure 2 gives both axes.

**Fig. 2.**
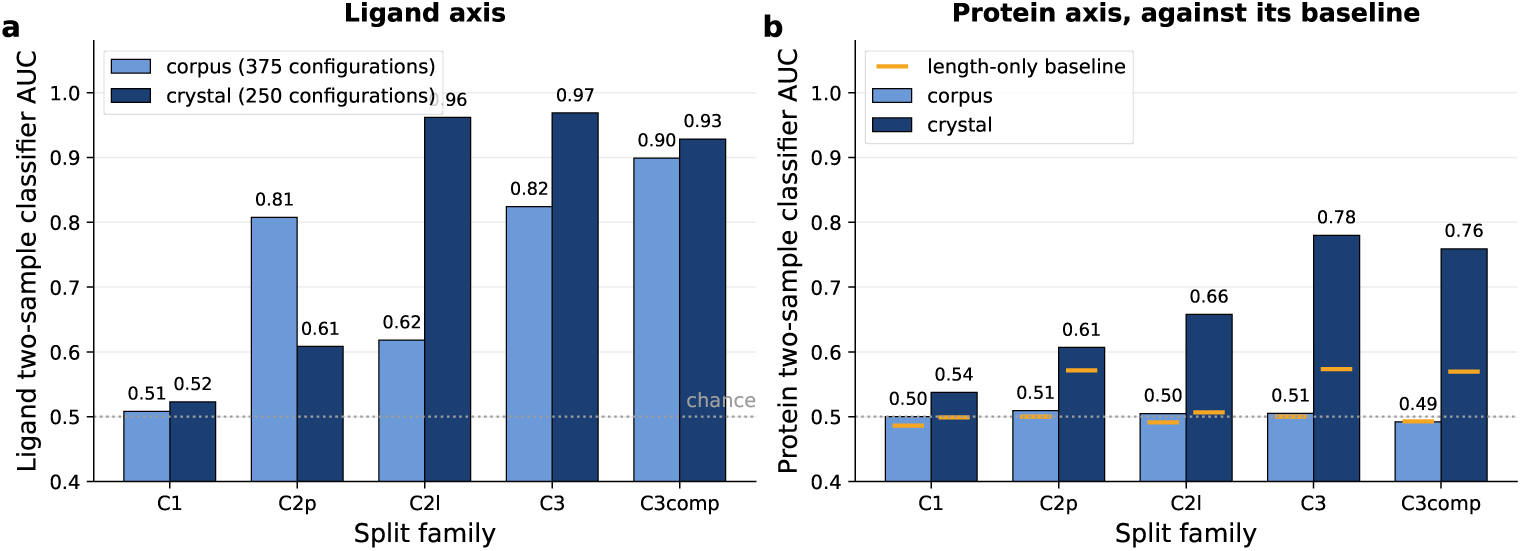
Distributional separation of the shipped split configurations, as the mean two-sample classifier AUC per family over 375 corpus and 250 crystal configurations. A score above the baseline means the held-out entities are distinguishable from the training entities, which is what a cold split is constructed to achieve. (a) Ligand axis, with chance marked. The two tiers behave oppositely, the crystal tier reaching 0.962 where the ligand is held out against 0.618 for the same family on the corpus. (b) Protein axis, each tier against the score the same discriminator reaches on protein length alone, which is 0.49 to 0.50 on the corpus and 0.5735 on the crystal tier. The crystal tier exceeds its baseline wherever an axis is held out and most strongly where both are, while every corpus family tracks its own baseline to within a percentage point, so partitioning corpus proteins does not by itself separate them beyond size.

Cross-fold sequence identity is reported under a coverage-normalised definition, the local identity scaled by alignment length over the shorter sequence, and the choice is measured rather than argued. Raw local identity is exactly 1.00 in every configuration of both tiers, since a seven-residue perfect match is reported as complete identity regardless of coverage. It therefore cannot distinguish any configuration from any other. The two measures differ by a factor of 75 on the crystal tier, where 9,455,750 ordered pairs exceed 30% under the raw definition against 125,832 under the normalised one. The coverage-normalised figure is meaningful only where the protein axis is held out, since a warm-protein family carries the same cluster on both sides by construction. Each configuration reports its own maximum, and across the protein-cold families those maxima run 0.2370 to 0.4780 on the corpus and 0.3740 to 0.9540 on the crystal tier. The two tiers are not comparable here. The lowest maximum of any protein-cold crystal configuration is 0.8410, so every one of them contains a cross-fold pair at least that similar, because the crystal tier is drawn from a structural archive dense with homologues and clustering at 30% identity leaves near-duplicates straddling the boundary. The guarantee is therefore redundancy reduction at a stated maximum rather than the absence of leakage, and the maximum is higher on the crystal tier than a user coming from the corpus would expect. Ligand-side separation is reported as the nearest train-neighbour Tanimoto, which runs 0.63 to 0.72 in the warm family and 0.31 to 0.39 in the jointly cold one.

One structural property bounds what the cold families can achieve. Neither entity axis contains a hub, since the largest protein cluster spans 2.0% of systems and the largest ligand cluster at most 1.1%. The bipartite system graph nonetheless percolates, and a single component holds between 77% and 99% of systems depending on the ligand cutoff. Hub-freeness and small components are different properties, and only the second bounds fold assignment.

#### Physical validity

Ligand poses were checked with PoseBusters[44] under the core validity criteria. Two restrictions apply and they are different in kind. By choice, the checks requiring a reference ligand in a shared frame were excluded, since no such frame is shared across the teachers. By availability, checking requires a reference ligand written in a form the harness can read, which exists for 13,391 of the 19,350 crystal-tier systems, of which 13,365 were scored, the remainder failing because no model file was present. For the cofolding arms the per-arm counts in Fig. 3 are the intersection of that reference set with the arm’s own coverage and are smaller than either. The docking arm’s denominators come from check resolution and are given in the figure caption. No rate below is computed on all 19,350 crystal systems, and none reaches unity even for experimental ligands, because the criteria impose an interatomic separation floor set by van der Waals radii on contacts that are not van der Waals contacts.

**Fig. 3.**
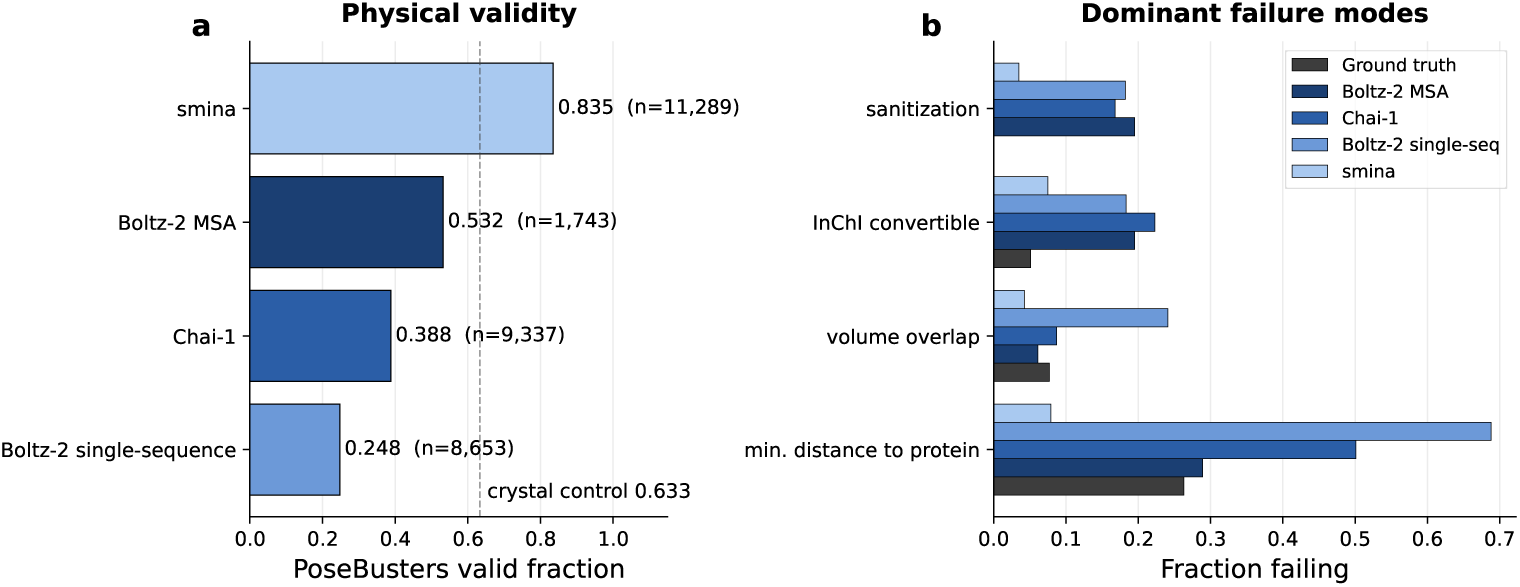
Physical validity under PoseBusters core criteria. (a) Valid fraction per arm with the experimental control at 0.633 marked, labelled Ground truth in panel b. The docking arm is scored under deposited-receptor conditioning to match it. (b) Fraction failing each of the four dominant checks. The checks do not share a denominator. Of the 11,290 docking poses, the two perception checks resolved on 11,289 and the two geometry checks on 11,195, the remainder carrying no conditioning receptor. Minimum distance to protein dominates on every arm, for the reason given in the text.

The 5,959 systems without a readable reference are not a random subset. They are dominated by metal cofactors and covalently attached ligands, the same classes that fail the distance check where they are present. The rates below are therefore computed on a set already depleted of the chemistry that drives the shortfall, and the control on the full tier would be lower.

The crystal tier provides the control that makes the rates interpretable, and that control is 0.633. The shortfall is concentrated rather than diffuse. Of the 990 experimental metal macrocycles in the crystal tier two pass, while on the remaining 12,375 systems the control rises to 0.683, and within that remainder the residual failures concentrate in covalently attached ligands and in phosphate-bearing cofactors. One check accounts for most of it. Minimum distance to protein fails on 0.263 of experimental ligands, and the classes it fails on are those whose contacts are coordination bonds, covalent linkages or salt bridges. A curated drug-like ligand set does not contain this chemistry, which is why published validity rates computed on such sets are not directly comparable to these.

Against that reference the teachers give 0.532 for the alignment-conditioned cofolder, 0.388 for Chai-1 and 0.248 for the single-sequence cofolder (Fig. 3). Two checks separate the cofolders from the control sharply. InChI convertibility and sanitisation fail on 16.8 to 19.5% of ligands from the three cofolding arms and on no experimental ligand at all, while the docking arm fails them on 7.5 and 3.5% respectively. Experimental ligands are deposited with a chemical component definition, whereas a predicted pose carries only the coordinates the model wrote, and the perception step that recovers bonds and valences from those coordinates is where the failures arise. These two checks therefore record a property of predicted output rather than a limitation of the checking software on this ligand set.

The ordering is the same on the full tier and on the non-macrocyclic subset of it, so the comparison does not depend on the ligand chemistry each arm happens to cover. On that subset the control is 0.683 and the arms are 0.542, 0.420 and 0.253, with the docking arm at 0.879. Every rate rises and none changes rank, which is what a shared cause predicts.

The docking arm is scored against the deposited receptor rather than against the one it was docked into, which puts it on the same conditioning as the crystal control and lets the two rates be read on one axis. Its poses are already in the crystal frame, so that conditioning is available to it and not to the cofolders. It reaches 0.835, above the 0.633 of the experimental control. That follows from the mechanism above. A docking engine samples against a scoring function that penalises short contacts, so it never enters the regime where the distance check misfires, while an experimental structure records that geometry wherever the chemistry produces it. Its internal geometry satisfies the bond and ring criteria by construction, as Methods describes.

These checks measure agreement with a van der Waals geometry model rather than correctness, and the control is what makes that visible.

#### Accuracy against ground truth

Two measures are reported together because they order the teachers differently and either alone would mislead. Symmetry-corrected ligand RMSD after pocket superposition measures where the ligand was placed. Contact-set agreement with the ground-truth label measures how much of the interaction the labels recover. The two were computed on different cohorts, the deviations during construction over a system list containing the deposited arms as a subset and the agreement on the deposited arms alone, so the counts in Fig. 4 exceed those in Table 7. These values are not comparable to the figures the teachers report on their own benchmarks, and the difference is configuration rather than evaluation. Chai-1 reports 77% of poses within 2 Å on the PoseBusters set against 32.1% here. Three differences account for the gap, namely one recycling step rather than the published setting, ESM embeddings in place of a multiple sequence alignment, and root-mean-square deviation computed after superposition on the pocket residues rather than under the benchmark protocol. A fourth applies to the crystal tier alone. 71.0% of its systems predate the Chai-1 training cutoff, so part of the published benchmark’s advantage and part of this table’s is recall. The configurations were chosen to reach corpus scale. What the table measures is those settings on these systems, and it should not be read as a benchmark of either method. Table 7 gives both per teacher and Fig. 4 shows the distributions.

**Fig. 4.**
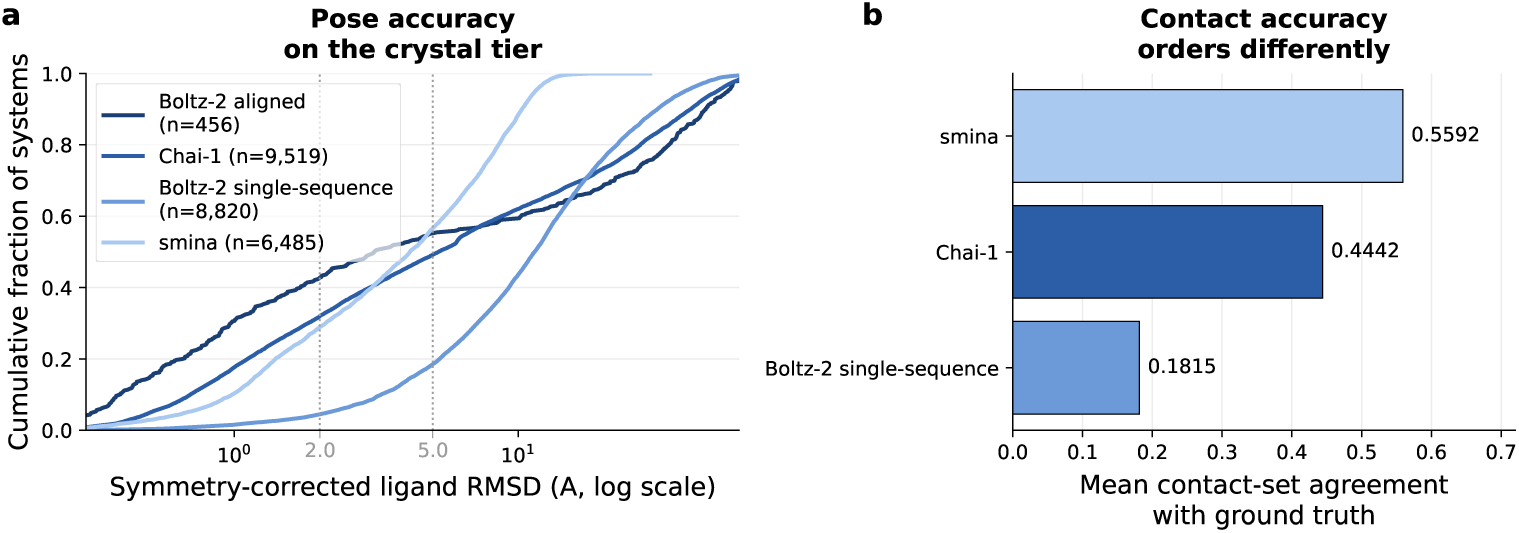
Two accuracy measures that order the teachers differently. (a) Cumulative distribution of symmetry-corrected ligand RMSD after superposition on the pocket residues, with the 2 and 5 Å marks indicated. For the cofolding arms the panel covers every resolved deviation, computed during construction over a system list that contains the deposited arm as a subset, so those medians differ in the second decimal from the Scored column of Table 7. The gap to that column is 150 systems on each cofolding arm, being the sum of two filters, deposition and then the requirement to carry distance labels, which coincide at the same total on both. The docking arm is shown on the paired subset that column also uses. (b) Mean contact-set agreement with the ground-truth label at 4 Å. Chai-1 places more ligands within 2 Å than the docking arm while recovering fewer contact residues and carrying a worse median deviation, so a user selecting on one measure should know what the other reports.

**Table 7.**
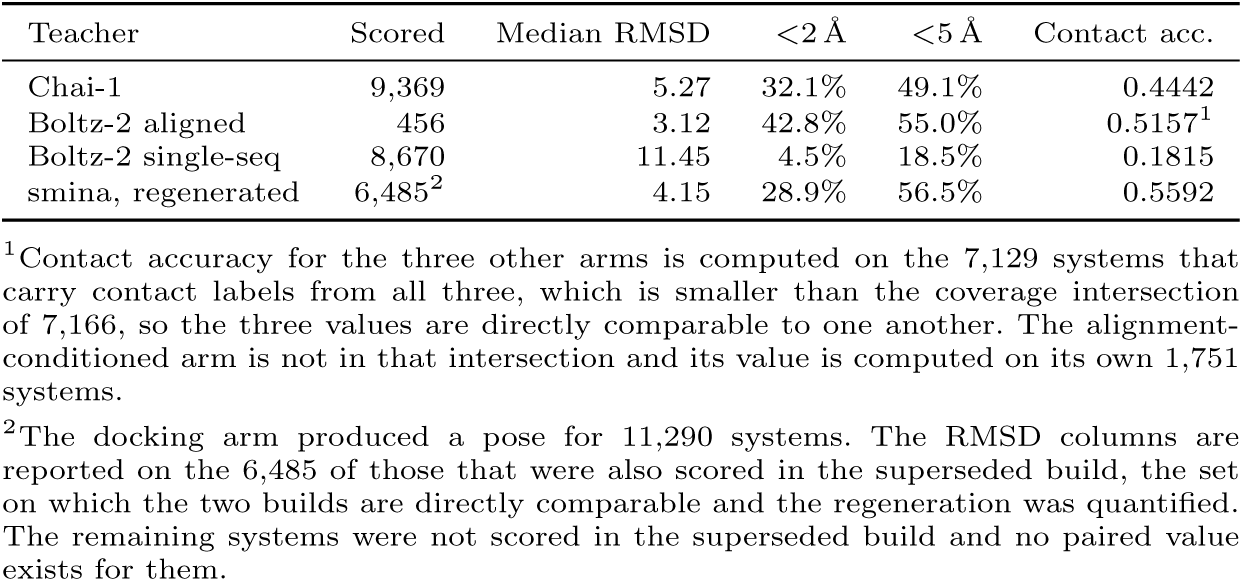
Accuracy against experimental ground truth on the crystal tier, reported to characterise the deposited labels and not to benchmark the methods against one another. RMSD is symmetry-corrected and computed after superposition on the pocket residues. Contact accuracy is the Jaccard overlap of the predicted and ground-truth contact residue sets at 4 Å. The two columns have different denominators, given in the footnotes.

The crystal tier is drawn from the same archive the cofolders were trained on, so part of the accuracy above is recall rather than prediction. Both training cutoffs are stated by their authors, 12 January 2021 for Chai-1 and 1 June 2023 for Boltz-2. 71.0% of crystal-tier systems were deposited before the first and 87.2% before the second, leaving 2,479 systems, 12.8% of the tier, which postdate both. Splitting the deposition date at each arm’s own cutoff bounds the effect. The scored cohort is poorer in recent structures than the tier it is drawn from, 18.5% of it postdating the Chai-1 cutoff against 29.0% of the full tier, because an older entry is more likely to carry a ligand the scoring harness can resolve. The comparison therefore reads the effect on less of the material where it would appear. Chai-1 places 33.4% of pre-cutoff ligands within 2 Å against 26.6% post-cutoff, with the median deviation rising from 4.98 to 6.15 Å. The single-sequence arm moves from 4.6% to 3.1%, a difference within the noise of a 550-system cohort. The effect is therefore present and bounded rather than absent, and the deposited rates should be read as an upper bound on what these configurations achieve on targets they have not seen. Two limits apply to the split itself. The postcutoff cohorts are 1,723 and 550 systems, and the source snapshot ends in early 2025, so the post-Boltz-2 window is under two years. The docking arm has no training cutoff, since it optimises a scoring function rather than learning from structures. That function was parameterised on protein–ligand complexes from the same archive, which is a weaker form of the same exposure and is not separable by deposition date.

Chai-1 against the docking arm is the clearest case of the two disagreeing. Chai-1 places the larger fraction within 2 Å, 32.1% against 28.9%, while the docking arm carries the lower median deviation, 4.15 against 5.27 Å, and recovers more contact residues, 0.5592 against 0.4442. A ligand can occupy the correct pocket in the wrong orientation, and the two measures are sensitive to different halves of that. The single-sequence arm is excluded from this comparison, since at 0.1815 contact accuracy and 4.5% within 2 Å it fails on both measures rather than disagreeing between them.

The regeneration is checked against the build it replaces on the systems both scored. Median RMSD falls from 8.80 to 4.15 Å and the fraction within 2 Å rises from 0.1% to 28.9%. This is the check on the regeneration described in Methods.

Contact-set agreement is not uniformly readable as accuracy. Where a teacher fails to reproduce the pocket backbone, the residue indices it reports can coincide with the ground-truth set without the pose being correct, and on constructs with repeated chains that coincidence is more likely. The affected systems are identifiable: they are those with a large pocket superposition residual, and the deposit records that residual per system so a user can exclude them. Agreement should be read alongside it rather than alone.

#### Reliability calibration

The per-system reliability annotation is an isotonic regression of observed label accuracy on inter-teacher agreement, constructed as Methods describes. Two error figures are reported for it and both compare the same two predictors, the isotonic map against a constant equal to the tier mean. Five-fold cross-validated, the map reaches a mean absolute error of 0.0814 against 0.1734 for the constant, so the estimate carries per-system information rather than reproducing that constant. On the single calibration split deposited with the fitted arrays and shown in Figure 5, the same comparison gives 0.0805 against 0.1752, so the fit does not depend on which half received it.

**Fig. 5.**
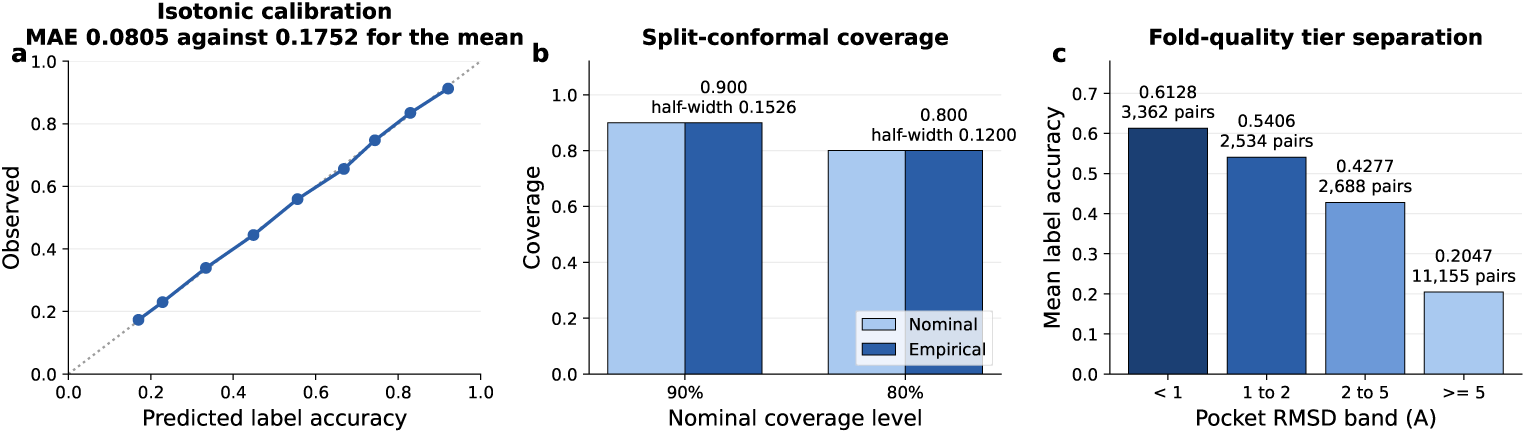
The reliability annotation, fitted on the crystal tier. (a) Isotonic calibration of observed label accuracy against the predicted value on the deposited calibration split, reaching a held-out mean absolute error of 0.0805 against 0.1752 for predicting the tier mean. (b) Split-conformal coverage, empirical against nominal, with half-widths 0.1526 at 90% and 0.1200 at 80%. The match is by construction and is shown as a check that the interval was built as Methods describes, not as independent evidence. (c) Mean label accuracy by pocket superposition residual band, over every crystal-tier teacher-system pair rather than the three-teacher core, so the band means do not average to the unfiltered figure in Table 8. The gradient shows that label accuracy tracks whether the pocket backbone was reproduced.

**Table 8.** Mean observed label accuracy against reliability threshold on the crystal tier, with the fraction retained. Observed accuracy is the mean contact-set agreement of the three covering teachers with the ground-truth label. The crystal column is of the 7,129 systems on which observed accuracy can be computed, and the corpus column of the systems carrying a reliability annotation, since no observed value exists there. Two cohort qualifications apply and neither is material. The crystal annotation cohort is 7,134 systems, five more than the observed cohort, and normalising on it would move no row by a tenth of a percentage point. The corpus percentages are computed on a base slightly smaller than the 23,433 systems the field covers, so a reader recomputing from the field will find each row low by about a tenth of a point.

| Threshold | Crystal kept | Crystal % | Mean accuracy | Corpus % |
| --- | --- | --- | --- | --- |
| none | 7,129 | 100.0 | 0.3951 | 100.0 |
| 0.20 | 6,081 | 85.3 | 0.4337 | 68.7 |
| 0.30 | 4,703 | 66.0 | 0.4938 | 35.3 |
| 0.40 | 3,495 | 49.0 | 0.5487 | 18.6 |
| 0.50 | 1,958 | 27.5 | 0.6287 | 8.6 |
| 0.60 | 823 | 11.5 | 0.7253 | 2.4 |
| 0.70 | 406 | 5.7 | 0.7924 | 0.5 |

The field is usable as a filter. Table 8 gives mean observed label accuracy among the crystal-tier systems retained at a series of thresholds, together with the fraction of each tier that survives. Accuracy rises monotonically from 0.3951 unfiltered to 0.7924 at a threshold of 0.70, at which point 5.7% of the crystal tier remains. The corpus tier retains a smaller fraction at every threshold, which is the covariate shift stated below appearing as a shift in the predicted distribution rather than as a failure of the ordering.

Intervals are split-conformal. Half-widths are 0.1526 at nominal 90% coverage and 0.1200 at 80%, and empirical coverage matches nominal at both. Two limits apply. The guarantee is marginal and not conditional, holding across the calibration distribution and not for any nominated system. And it holds on the crystal tier alone, because split-conformal coverage assumes the calibration and target distributions are exchangeable[55]. They are not across tiers. Mean agreement is 0.225 on the crystal tier against 0.108 on the corpus tier, and the density ratio between the two spans roughly two orders of magnitude across the agreement range, with corpus systems concentrated at the low end where the crystal calibration set is sparsest. Recovering coverage under a shift of this kind requires weighting the calibration residuals by an estimated likelihood ratio between the two distributions[56], and that construction is not applied here. The deposited half-width is therefore to be read on the crystal tier alone, and pred accuracy on the corpus tier as a calibrated ordering and not a certified accuracy.

Per-residue support is the direct test of whether multi-teacher agreement buys anything, and it does. Precision against the ground-truth contact set rises from 0.2175 where one teacher reports a residue, to 0.7265 at two and 0.9356 at three, computed over the full crystal-tier support table rather than a sample, while the union of all three recovers 0.8828 of ground-truth contact residues (Fig. 6). The distribution of that count is uneven. 70.7% of crystal residue rows carry a single asserting teacher and 7.7% are unanimous, so filtering to unanimous contacts buys precision at the cost of most of the rows and inherits whatever composition the agreeing subset has. That is a property of the labels rather than of any model trained on them, and it is what the reliability annotation aggregates to the system level. A demonstration that a model trained on these labels outperforms one trained on single-teacher labels is deliberately not attempted here, since this article describes the resource and a training study is a separate contribution with its own controls.

**Fig. 6.**
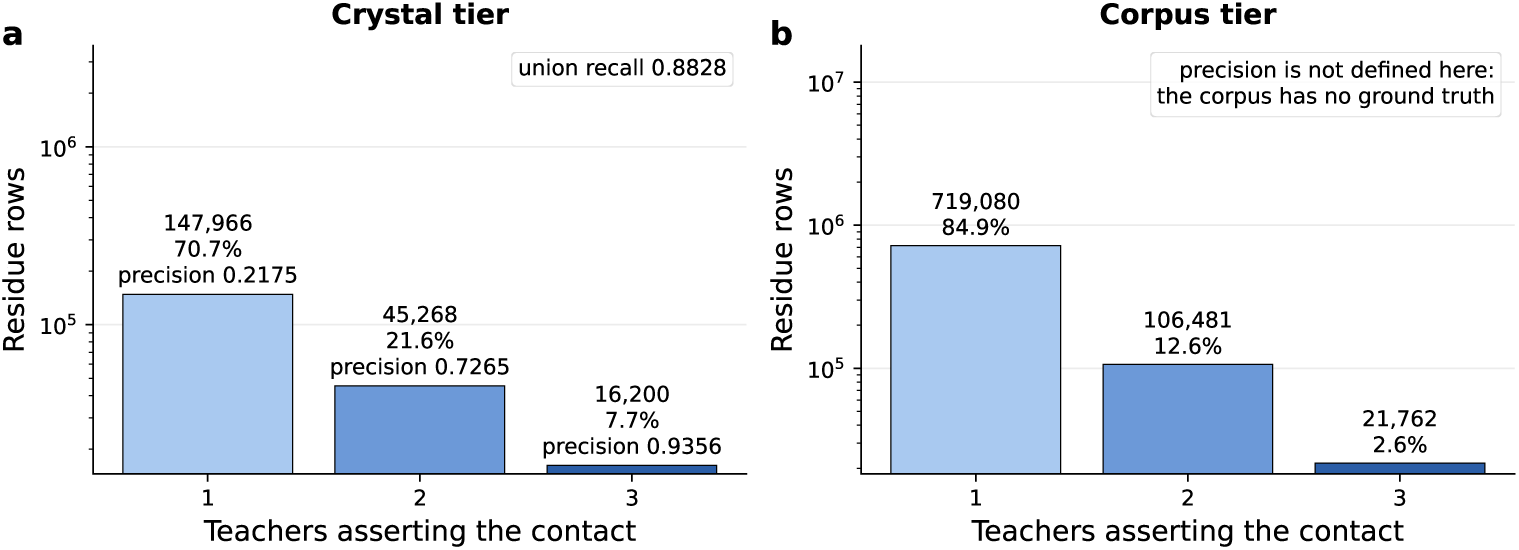
Per-residue teacher support. (a) Crystal tier, where precision against the ground-truth contact set rises from 0.2175 at one asserting teacher to 0.7265 at two and 0.9356 at three, while the union recovers 0.8828 of ground-truth contact residues. (b) Corpus tier, where the same distribution is available but precision is undefined because no experimental complex exists. Row counts are residue rows, not systems.

Applied to the corpus the estimate gives a mean predicted accuracy of 0.2784 against 0.3947 on the crystal tier it was fitted on, a difference consistent with the corpus occupying the low-agreement region of a crystal-fitted scale. On the crystal tier the mean predicted value matches the mean observed value of 0.3951 to within 0.0004, which is what a correctly fitted isotonic map gives and is not evidence about the corpus. Corpus labels should be read as predictions carrying a calibrated reliability ordering rather than as measurements, in the sense that predicted structures are hypotheses rather than observations[53].

#### Reproducibility

Regenerating a prediction does not return the same coordinates. Re-running the single-sequence cofolder on byte-identical inputs under the recorded seed returns structures that differ from the deposited ones, and the difference grows with system size. The deposited coordinates are therefore one draw from the configuration recorded in Table 2 rather than a reproducible artifact, and a user regenerating them should expect agreement in aggregate and not per system. The seed is not the only dependence. The arms were produced under the versions given in the Software subsection, and a later version of the same predictor does not reproduce them either. This is a documented property of the model family rather than of this pipeline. A published evaluation of cofolding on a related docking task found variation across seeds large enough that stable analysis required sampling an ensemble rather than a single seed[57].

Each arm is one configuration sampled once, not an ensemble. What the deposit substitutes for that is disagreement across methods of different construction, which samples a wider space than reseeding one method does, since two architectures fail differently where one architecture fails twice the same way. It is not a replacement. The reliability annotation describes disagreement between teachers at a fixed configuration and is a lower bound on the variation a user would see across repeated runs of any one of them.

#### Usage Notes

Which layer to read follows from the task. Labels at the deposited cutoff come from the distance tables, and any other cutoff, representation or pair definition from the coordinate store. Within the store each protein representation is a column selection rather than a conversion, and Table 5 gives the views and their derivations, each exposed by a converter in the code release.

Reduced views are reduced. A side-chain centroid fixes the direction and extent of a side chain and not its rotamer state, and the published reconstruction accuracies for such representations[36] are those of a trained network rather than of direct geometric inversion. The deposited labels are distances and not interaction types, so hydrogenbond geometry, stacking and ionic contacts are not annotated. The shell coordinates support deriving them, and a worked example ships with the deposit as a notebook. A distance recomputed from two stored coordinates typically differs from the deposited value by about 0.004 Å, twice the verified per-coordinate round trip, and by at most about 0.02 Å, since each coordinate can sit half a quantisation step from its true position. The bound is the margin to allow when a recomputed distance is compared against a threshold. The population at risk is small but not negligible. 22,960 rows sit exactly on the 4 Å boundary, 37,713 on the 5 Å boundary and 128,932 on the 8 Å boundary, so a flag recomputed at a different precision can differ from the deposited one on those rows alone.

A user training a model that takes three-dimensional structure at input and derives distance maps in the forward pass reads the coordinate store and ignores the distance tables. Ligand-to-residue, residue-to-residue and ligand-to-ligand maps are all computed from the stored coordinates at whatever cutoff and representation the model uses, and only the first of the three is present in the tables at any cutoff. The tiers divide naturally, and the division allows one construction in particular. Each configuration’s output is stored separately on the teacher axis, so a model could carry one prediction head per teacher, trained to reproduce what each configuration produces for a given pair including where it is wrong, with a further head fitted to experiment where experiment exists and to the teachers’ consensus weighted by the agreement value where it does not. Whether such a model outperforms one fitted to the crystal tier alone is an open question this deposit is built to let someone answer, and not a claim it makes.

That division has a known cost and the deposit does not evade it. Placing a ligand against a predicted receptor is a documented failure mode, and the corpus tier is that case on every one of its systems. AlphaFold2 models reproduce binding-pocket geometry well, but docked pose accuracy into those models does not improve correspondingly and falls short of docking into experimental structures solved without the ligand present[58], a result reported for G protein-coupled receptors and corroborated across several independent evaluations. The corpus receptors are from a later generation of the same predictor. It is not exposed to the second documented failure mode in the same way, since a pair never co-crystallised together is not one a model can have reproduced from training[11], but the first alone is sufficient reason to treat corpus geometry as hypothesis. The reliability annotation is the response rather than a denial, calibrated where accuracy is measurable and applied where it is not, so a user filtering on pred accuracy is selecting against the systems where the failure mode bites hardest. The alternative without it would be to take the corpus tier whole or to discard it, neither of which distinguishes the systems the failure mode reaches from those it does not. Table 8 gives what the filtering costs in coverage.

A user evaluating what the teachers produce reads the crystal tier, where deposited coordinates anchor the predictions. Two properties bound what that evaluation measures. The teacher configurations are throughput-oriented, so the coordinates record what those settings produce and not what the methods achieve on a single target configured for accuracy. And the receptor conformations carry no reliable inducedfit signal outside the crystal tier. Docking holds the receptor rigid by construction, and cofolding models predict a receptor conformation but are reported not to reproduce the rearrangements a ligand induces[59], so ligand-adapted receptor geometry is present only where the experiment put it. The arms are also dated. They record the behaviour of the versions named in Table 2 and should not be read as the current state of either method.

646 split configurations are provided and the choice among them is deliberate. The warm family supports interpolation claims, while the joint-cold C3 and C3comp families support prospective ones and carry the lowest residual cross-fold identity of any family on either tier. The protein identity threshold should be selected against the claim being made rather than defaulted. A user wanting a smaller, higher-quality subset can filter on the reliability field, and Table 8 gives the trade, both of its columns being fractions of the reliability-annotated cohort rather than of the tier. At a threshold of 0.40 the crystal tier retains 3,495 of its 7,129 annotated systems at a mean label accuracy of 0.5487, and the corpus retains 4,326 of 23,433. Filtering the corpus to that label quality therefore leaves it the size of the crystal tier, so a user wanting both scale and quality does not have them simultaneously.

Five boundaries apply. First, the per-system reliability field is a calibrated ordering on the corpus rather than certified coverage, since the mapping is fitted on the crystal tier and does not transfer unweighted across that shift. Second, the three-teacher intersection over-represents small, non-metal ligands and contains an interface binder in 14.6% of systems against 32.4% outside it, so it should not carry a benchmark for large, metal-containing or interface systems. Third, the corpus protein set inherits a partition built to control protein-protein sequence leakage. What it guarantees is sequence non-redundancy and not pocket diversity, so a protein-cold split here demonstrates generalisation across sequence space and not across pocket space, and two accessions with unrelated sequences may still present similar binding sites. Fourth, the distance tables are censored rather than complete, since a crystal-tier system carries rows for 23.5% of its residues on average and the remainder lie beyond the 15.0 Å cutoff, so an outer join filling the absent pairs with zero records a contact where none was measured. Fifth, the deposited shell is defined by distance to the ligand and holds only the heavy atoms of standard residues, so a model given the shell alone has been told where to look and is not seeing the metal ions or cofactors inside the cutoff. A user who needs a pocket that does not depend on the answer takes the corpus pocket annotation, which the crystal tier does not have, or locates one on the stored C*α* trace.

## Data Availability

The dataset is deposited at Zenodo under the version DOI 10.5281/zenodo.21560088[48] and released under CC BY 4.0. The concept DOI 10.5281/zenodo.21560087 resolves to the most recent version. A convenience mirror is maintained at https://huggingface.co/datasets/ThorKl/PLI-parallax, under a Hugging Face account name that differs from the GitHub one. The Zenodo record is the canonical release, and the mirror carries the same bytes, so the checksums recorded in the deposit manifest verify against either copy. Upstream sources are listed with their versions or access dates in Table 3. One of them, ChEMBL, is share-alike licensed, and no ChEMBL-derived value reaches the deposit. The activity threshold determined which pairs were assembled and no activity value is stored, while the deposited ligand structures are keyed on InChIKey and system identifier and carry no ChEMBL accession, so the share-alike term does not propagate to the release licence. The deposit contains predicted coordinates and derived distance labels rather than newly determined experimental structures, so it carries no depositions for a structural repository. All coordinates in the crystal tier were read from existing Protein Data Bank entries[26] and are not redistributed as such, and no field value from any upstream annotation resource appears in the deposit.

## Code Availability

The code that produced the deposit is at https://github.com/ThorKlm/PLI-parallax under Apache 2.0, archived at Zenodo under 10.5281/zenodo.22258211, with the tagged release v1.0.0 the state of the code as it was run. It carries the validation suite of 57 checks and verify quoted figures v2.py, which recomputes 130 of the figures quoted here directly from the deposited Parquet and names the thirteen it does not reach. Both take the deposit root as an argument. A usage notebook ships with the deposit and in the repository under notebooks/. It fetches the deposit, verifies it against MANIFEST.sha256, recomputes a distance table from the coordinate store and draws the three-bead representation across teachers, and it runs in Colab without a local installation.

Three stages are not reproducible from the repository alone. The conversion from intermediate prediction output to the deposited tables is performed by scripts that are archived but not tied into a single driver. The corpus pocket annotation was produced by a detector ensemble maintained separately and is deposited as a finished artifact. The reliability fit consumed an intermediate agreement table that is not deposited, so the fit cannot be repeated exactly as it was run. The annotation itself is nonetheless re-derivable: the agreement column recomputes from the deposited contact tables to the fourth decimal, and the fitted arrays ship alongside the field, so a reader can reconstruct pred accuracy from what the deposit contains. Software versions are given in the Software subsection of the Methods.

## Author contributions

T.K.: Conceptualization, Methodology, Software, Validation, Formal analysis, Investigation, Data curation, Writing – original draft, Visualization, Project administration.

## Ethics approval

Not applicable. The work uses only publicly deposited structural and bioactivity data and involves no human or animal subjects.

## Acknowledgements

The author thanks the maintainers of the upstream resources listed in Table 3 for making their data openly available, and the developers of the prediction and analysis tools cited in the Software subsection.

## References

[1] Nittinger, E., Yoluk, Ö., Tibo, A., Olanders, G. & Tyrchan, C. Co-folding, the future of docking – prediction of allosteric and orthosteric ligands. Artificial Intelligence in the Life Sciences 8, 100136 (2025).

[2] Wan, S., Zhang, X., Xue, X. & Coveney, P. V. Reliability of AI methods in drug discovery: evaluation of Boltz-2 for structure and binding affinity prediction. Journal of Chemical Theory and Computation 22, 7811–7824 (2026).

[3] Wang, R., Fang, X., Lu, Y. & Wang, S. The PDBbind database: Collection of binding affinities for protein-ligand complexes with known three-dimensional structures. Journal of Medicinal Chemistry 47, 2977–2980 (2004).

[4] Wang, Y. et al. A workflow to create a high-quality protein-ligand binding dataset for training, validation, and prediction tasks. Digital Discovery 4, 1209–1220 (2025).

[5] Li, J. et al. Leak proof PDBBind: A reorganized data set of protein-ligand complexes for more generalizable binding affinity prediction. The Journal of Physical Chemistry B 130, 730–740 (2026).

[6] Durairaj, J. et al. PLINDER: The protein-ligand interactions dataset and evaluation resource (2024). Preprint at 10.1101/2024.07.17.603955.

[7] Siebenmorgen, T., et al. MISATO: machine learning dataset of protein-ligand complexes for structure-based drug discovery. Nature Computational Science 4, 367–378 (2024).

[8] Lemos, P. et al. SAIR: Enabling deep learning for protein-ligand interactions with a synthetic structural dataset (2025). Preprint at 10.1101/2025.06.17.660168.

[9] Wohlwend, J. et al. Boltz-1: democratizing biomolecular interaction modeling (2025). Preprint at 10.1101/2024.11.19.624167.

[10] Wei, J. et al. GatorAffinity: boosting protein-ligand binding affinity prediction with large-scale synthetic structural data (2025). Preprint at 10.1101/2025.09.29.679384.

[11] Škrinjar, P., et al. Evaluating generalization in protein-ligand cofolding methods. Nature Structural & Molecular Biology 33, 782–794 (2026).

[12] Hetmann, M. et al. Folding the human proteome using BioNeMo: a fused dataset of structural models for machine learning purposes. Scientific Data 11, 591 (2024).

[13] Zhang, Z. et al. Structure prediction of novel isoforms from uveal melanoma by AlphaFold. Scientific Data 10, 513 (2023).

[14] Ektefaie, Y. et al. Evaluating generalizability of artificial intelligence models for molecular datasets. Nature Machine Intelligence 6, 1512–1524 (2024).

[15] Shihab, I. F., Akter, S. & Sharma, A. CalPro: prior-aware evidential-conformal prediction with structure-aware guarantees for protein structures. arXiv (2026). Preprint.

[16] Hong, Y., et al. Trustworthy protein-ligand binding affinity prediction via reliability-aware multi-engine fusion, 694–699 (ACM, 2026).

[17] Francoeur, P. G. et al. Three-dimensional convolutional neural networks and a cross-docked data set for structure-based drug design. Journal of Chemical Information and Modeling 60, 4200–4215 (2020).

[18] Houston, D. R. & Walkinshaw, M. D. Consensus docking: improving the reliability of docking in a virtual screening context. Journal of Chemical Information and Modeling 53, 384–390 (2013).

[19] Chai Discovery team et al. Chai-1: Decoding the molecular interactions of life (2024). Preprint at 10.1101/2024.10.10.615955.

[20] Passaro, S. et al. Boltz-2: Towards accurate and efficient binding affinity prediction (2025). Preprint at 10.1101/2025.06.14.659707.

[21] Abramson, J. et al. Accurate structure prediction of biomolecular interactions with AlphaFold 3. Nature 630, 493–500 (2024).

[22] Koes, D. R., Baumgartner, M. P. & Camacho, C. J. Lessons learned in empirical scoring with smina from the CSAR 2011 benchmarking exercise. Journal of Chemical Information and Modeling 53, 1893–1904 (2013).

[23] Jumper, J. et al. Highly accurate protein structure prediction with AlphaFold. Nature 596, 583–589 (2021).

[24] Bertoni, D. et al. AlphaFold protein structure database 2025: a redesigned interface and updated structural coverage. Nucleic Acids Research 54, D358–D362 (2026).

[25] Zhang, C., Zhang, X., Freddolino, L. & Zhang, Y. BioLiP2: an updated structure database for biologically relevant ligand-protein interactions. Nucleic Acids Research 52, D404–D412 (2024).

[26] Berman, H. M. et al. The Protein Data Bank. Nucleic Acids Research 28, 235–242 (2000).

[27] Bernett, J., Blumenthal, D. B. & List, M. Cracking the black box of deep sequence-based protein-protein interaction prediction. Briefings in Bioinformatics 25, bbae076 (2024).

[28] Liu, T. et al. BindingDB in 2024: a FAIR knowledgebase of protein-small molecule binding data. Nucleic Acids Research 53, D1633–D1644 (2025).

[29] Zdrazil, B. et al. The ChEMBL database in 2023: a drug discovery platform spanning multiple bioactivity data types and time periods. Nucleic Acids Research 52, D1180–D1192 (2024).

[30] Mirdita, M. et al. ColabFold: making protein folding accessible to all. Nature Methods 19, 679–682 (2022).

[31] ByteDance AML AI4Science Team et al. Protenix – advancing structure prediction through a comprehensive AlphaFold3 reproduction (2025). Preprint at 10.1101/2025.01.08.631967.

[32] Park, Y. & Marcotte, E. M. Flaws in evaluation schemes for pair-input computational predictions. Nature Methods 9, 1134–1136 (2012).

[33] Steinegger, M. & Söding, J. MMseqs2 enables sensitive protein sequence searching for the analysis of massive data sets. Nature Biotechnology 35, 1026–1028 (2017).

[34] Bemis, G. W. & Murcko, M. A. The properties of known drugs. 1. molecular frameworks. Journal of Medicinal Chemistry 39, 2887–2893 (1996).

[35] Graber, D. et al. Resolving data bias improves generalization in binding affinity prediction. Nature Machine Intelligence 7, 1713–1725 (2025).

[36] Heo, L. & Feig, M. One bead per residue can describe all-atom protein structures. Structure 32, 97–111.e6 (2024).

[37] Rotkiewicz, P. & Skolnick, J. Fast procedure for reconstruction of full-atom protein models from reduced representations. Journal of Computational Chemistry 29, 1460–1465 (2008).

[38] Zadrozny, B. & Elkan, C. Transforming classifier scores into accurate multiclass probability estimates, 694–699 (ACM, 2002).

[39] Vovk, V., Gammerman, A. & Shafer, G. Algorithmic Learning in a Random World 2 edn (Springer, Cham, 2022).

[40] Norinder, U., Carlsson, L., Boyer, S. & Eklund, M. Introducing conformal prediction in predictive modeling. a transparent and flexible alternative to applicability domain determination. Journal of Chemical Information and Modeling 54, 1596–1603 (2014).

[41] O’Boyle, N. M. et al. Open Babel: An open chemical toolbox. Journal of Cheminformatics 3, 33 (2011).

[42] Landrum, G., et al. RDKit: open-source cheminformatics (2026). Release 2026.03.3, https://www.rdkit.org.

[43] Wojdyr, M. GEMMI: A library for structural biology. Journal of Open Source Software 7, 4200 (2022).

[44] Buttenschoen, M., Morris, G. M. & Deane, C. M. PoseBusters: AI-based docking methods fail to generate physically valid poses or generalise to novel sequences. Chemical Science 15, 3130–3139 (2024).

[45] Meli, R. & Biggin, P. C. spyrmsd: symmetry-corrected RMSD calculations in Python. Journal of Cheminformatics 12, 49 (2020).

[46] Pedregosa, F. et al. Scikit-learn: Machine learning in Python. Journal of Machine Learning Research 12, 2825–2830 (2011).

[47] Trott, O. & Olson, A. J. AutoDock Vina: Improving the speed and accuracy of docking with a new scoring function, efficient optimization, and multithreading. Journal of Computational Chemistry 31, 455–461 (2010).

[48] Klamt, T. PLI-Parallax: protein-ligand coordinates from multiple structure predictors and crystal structures with distance and contact labels (2026). Zenodo 10.5281/zenodo.21560088.

[49] The UniProt Consortium. UniProt: the Universal Protein Knowledgebase in 2025. Nucleic Acids Research 53, D609–D617 (2025).

[50] Le Guilloux, V., Schmidtke, P. & Tuffery, P. Fpocket: An open source platform for ligand pocket detection. BMC Bioinformatics 10, 168 (2009).

[51] Krivák, R. & Hoksza, D. P2Rank: machine learning based tool for rapid and accurate prediction of ligand binding sites from protein structure. Journal of Cheminformatics 10, 39 (2018).

[52] Sestak, F. et al. VN-EGNN: E(3)and SE(3)-equivariant graph neural networks with virtual nodes enhance protein binding site identification. Journal of Cheminformatics 18, 11 (2026).

[53] Terwilliger, T. C. et al. AlphaFold predictions are valuable hypotheses and accelerate but do not replace experimental structure determination. Nature Methods 21, 110–116 (2024).

[54] Lopez-Paz, D. & Oquab, M. Revisiting classifier two-sample tests (ICLR, 2017).

[55] Barber, R. F., Candés, E. J., Ramdas, A. & Tibshirani, R. J. Conformal prediction beyond exchangeability. The Annals of Statistics 51 (2023).

[56] Tibshirani, R. J., Barber, R. F., Candés, E. J. & Ramdas, A. Conformal prediction under covariate shift, Vol. 32, 2530–2540 (Curran Associates, 2019).

[57] Hitawala, F. N. & Gray, J. J. What does AlphaFold3 learn about antibody and nanobody docking, and what remains unsolved? mAbs 17, 2545601 (2025).

[58] Karelina, M., Noh, J. J. & Dror, R. O. How accurately can one predict drug binding modes using AlphaFold models? eLife 12, RP89386 (2023).

[59] Kim, J. et al. Large scale prospective evaluation of co-folding across 557 Mac1ligand complexes and three virtual screens (2025). Preprint at 10.64898/2025.12.25.696505.

